# Accuracy of somatic variant detection in multiregional tumor sequencing data

**DOI:** 10.1101/655605

**Authors:** Harald Detering, Laura Tomás, Tamara Prieto, David Posada

## Abstract

Multiregional bulk sequencing data is necessary to characterize intratumor genetic heterogeneity. Novel somatic variant calling approaches aim to address the particular characteristics of multiregional data, but it remains unclear to which extent they improve compared to single-sample strategies. Here we compared the performance of 16 single-nucleotide variant calling approaches on multiregional sequencing data under different scenarios with in-silico and real sequencing reads, including varying sequencing coverage and increasing levels of spatial clonal admixture. Under the conditions simulated, methods that use information across multiple samples do not necessarily perform better than some of the standard calling methods that work sample by sample. Nonetheless, our results indicate that under difficult conditions, Mutect2 in multisample mode, in combination with a correction step, seems to perform best. Our analysis provides data-driven guidance for users and developers of somatic variant calling tools.

## Introduction

Pervasive intratumor heterogeneity (ITH) has been reported in many cancer types, usually based on single tumor samples[1]. Deciphering ITH is essential for understanding cancer development[2] and inform treatment[3, 4]. However, there are inherent limitations to characterize ITH using single biopsies[5]. Importantly, subclonal mutations can easily remain undetected if they are not present at a minimal frequency in a given sample. Not surprisingly, early multiregional sequencing (M-seq) studies in different cancer types showed that a substantial fraction of the somatic mutations are exclusively seen in a subset of the samples[5–8]. Obviously, the analysis of multiple regional samples from the same tumor should in general increase the power to detect subclonal variants and therefore provide a more accurate description of ITH.

Historically, studies involving M-seq data detected somatic variants by independently analyzing single biopsies[9–11], i.e., following a marginal calling strategy. More recently, specific approaches have been proposed for the detection of somatic single nucleotide variants (sSNVs) from M-seq data. Some strategies first detect variants independently for each region, and in a second step filter them (i.e, two-step calling), like MuClone[12] or SNV-PPILP[13]. Alternatively, methods like MultiSNV[14] or HaplotypeCaller[15] model the presence of mutations across multiple samples at once, i.e., a joint calling strategy. Mutect2[16] now also offers a multisample mode, in beta stage.

Multiple studies have benchmarked somatic mutation callers using single biopsies[17–19], but only two have specifically evaluated somatic variant calling for multiregional data, in both cases when presenting a new calling method[12],[14]. To our knowledge, an independent assessment of variant callers using M-seq tumor data has never been carried out. Indeed, third-party benchmarking is important because authors evaluating their own methods against others may be prone to the “self-assessment trap”[20], i.e., implicit biases in the evaluation conditions. Besides, previous comparisons have explored a limited range of rather simple multiregional scenarios. In order to fill this gap, here we carried out a comprehensive comparison of 16 approaches for sSNV detection in M-seq data using two different simulation strategies, with in silico and real sequencing reads. Our results unveil the relative performance of different variant calling strategies when dealing with M-seq data.

## Results

We simulated 120 *de novo* and 30 *spike*-*in* tumors from which we obtained five multiregional samples (Fig. 1). For the former, we generated *in silico* sequencing reads at three clonal admixture levels and four sequencing depths. For the latter, we introduced mutations in real reads from a healthy sample at three clonal admixture levels.

**Figure 1:**
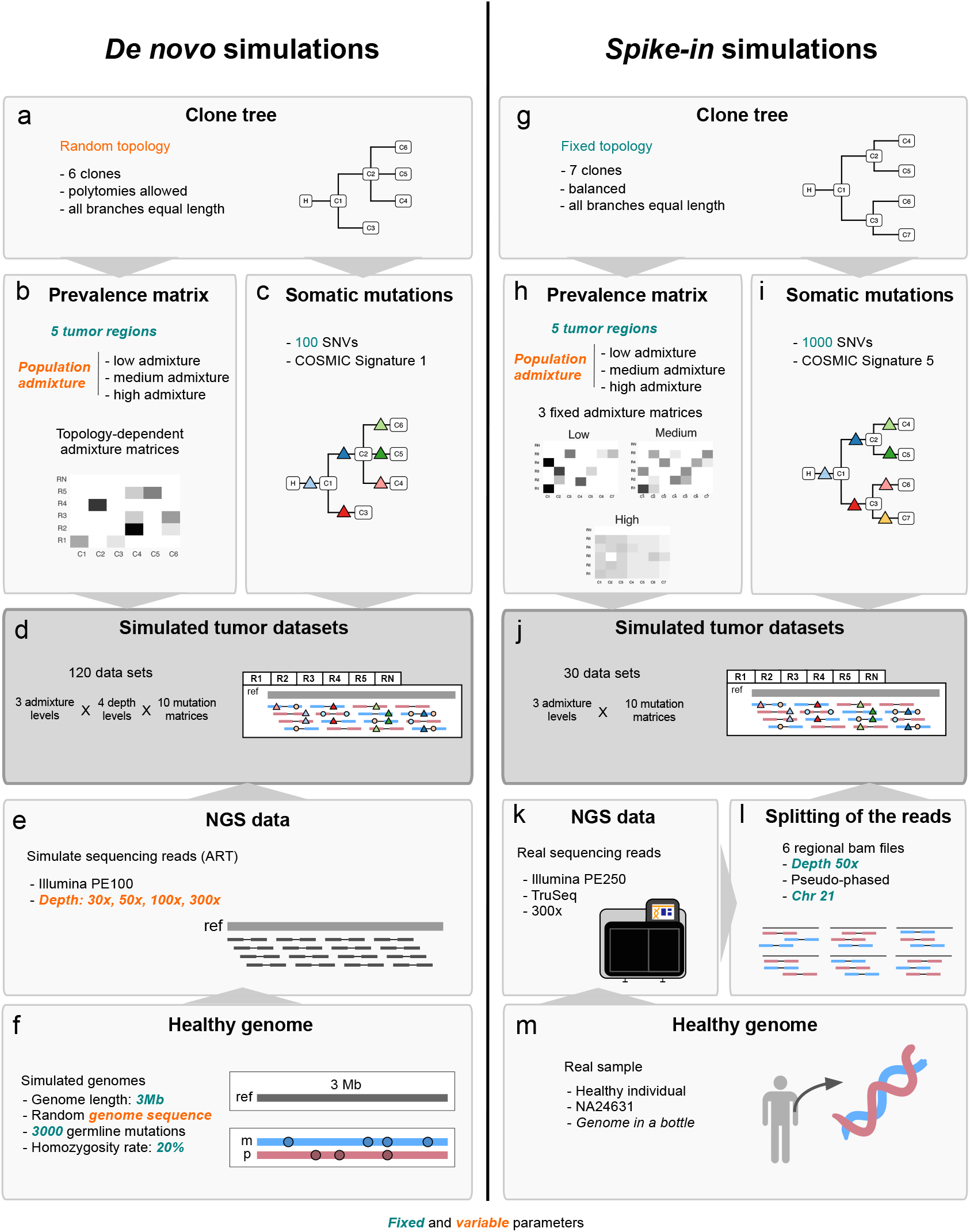
Simulation steps to generate *de novo* and *spike*-*in* simulated M-seq data sets. More details in Supplementary Information.

### Overall variant calling performance in the de novo simulation

In general, most variant callers showed median recall values above 0.6, while Shimmer and SNooPer at low depths, and VarScan, MuClone and particularly SNV-PPILP at all sequencing depths, performed worse (Fig. 2). Mutect2 in multisample mode plus the FP filter (Mutect2_multi_F, see “Variant filtering” section in Online Methods) achieved the highest recall at 30x and 50x, while perfect recall was obtained by Mutect2_multi_F and VarDict at 100x and 300x, as well as by Mutect2_single, Shimmer, Strelka2 at 300x. The recall rate of Bcftools, SomaticSniper, VarScan, MuClone and SNV-PPILP was not affected significantly by sequencing depth, while the remaining methods performed better at higher depths. Precision was generally very high, often with values above 0.9, but with some exceptions. SNooPer was consistently unprecise (< 0.2) independent of sequencing depth. Surprisingly, VarDict’s precision decreased dramatically with increasing depth. The precision of Mutect2_multi_F was also slightly lower at higher depths. As precision values were usually good, F1 scores mainly reflected differences in recall. Mutect2_multi_F achieved the highest F1 score at 30x and 50x, Mutect2_single scored best at 100x, while Mutect2_single, Shimmer and Strelka2 all achieved perfect performance at 300x. In general, Mutect2_multi_F, MultiSNV, Mutect2_single, MuTect1 and Strelka2 were best (pairwise Wilcoxon rank sum test P-value > 0.05).

**Figure 2:**
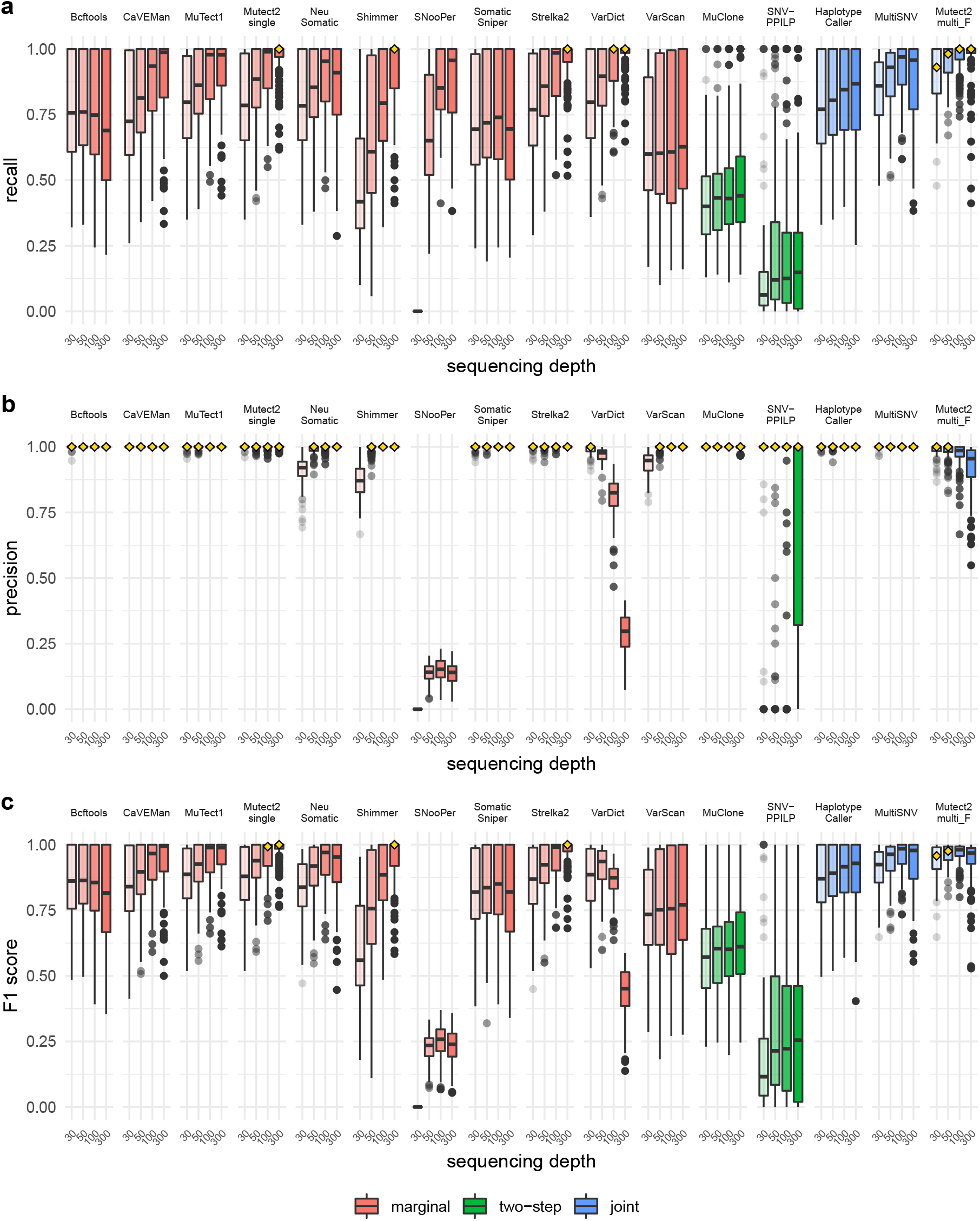
Performance of variant callers in the *de novo* simulation at different sequencing depths. **a** recall. **b** precision. **c** F1 score. In the boxplots the central line indicates the median, while the box limits correspond to the *Q*1 and *Q*3 quartiles; upper and lower whiskers extend from *Q*3 to *Q*3 + 1.5(*Q*3 − *Q*1) and from *Q*1 to *Q*1 − 1.5(*Q*3 − *Q*1), respectively. All metrics were calculated per tumor sample. The highest median score in each category is highlighted with a yellow diamond.

### Effect of clonal admixture in the de novo simulation

As a general trend, higher admixture was associated with lower recall, with the exception of Mutect2_multi_F, MuClone and SNV-PPILP (Fig. 3). The highest median recall in low admixture tumors was achieved by Bcftools, CaVEMan, MuTect1, Mutect2_single, NeuSo-matic, Shimmer, SomaticSniper, Strelka2, VarDict, HaplotypeCaller and Mutect2_multi_F. Mutect2_multi_F also accomplished the best recall in the medium and high admixture cases. Precision was independent of the admixture level. The trend for the F1 scores was the same as for recall, except for Mutect2_multi_F, MuClone, SNooPer and SNV-PPILP which were unaffected by admixture. While Bcftools, CaVEMan, MuTect1, Mutect2_single, SomaticSniper, Strelka2 and HaplotypeCaller showed optimal performance at low admixture, Mutect2_single achieved the top F1 score at medium admixture levels, while Mutect2_multi_F outperformed all other methods when admixture was high.

**Figure 3:**
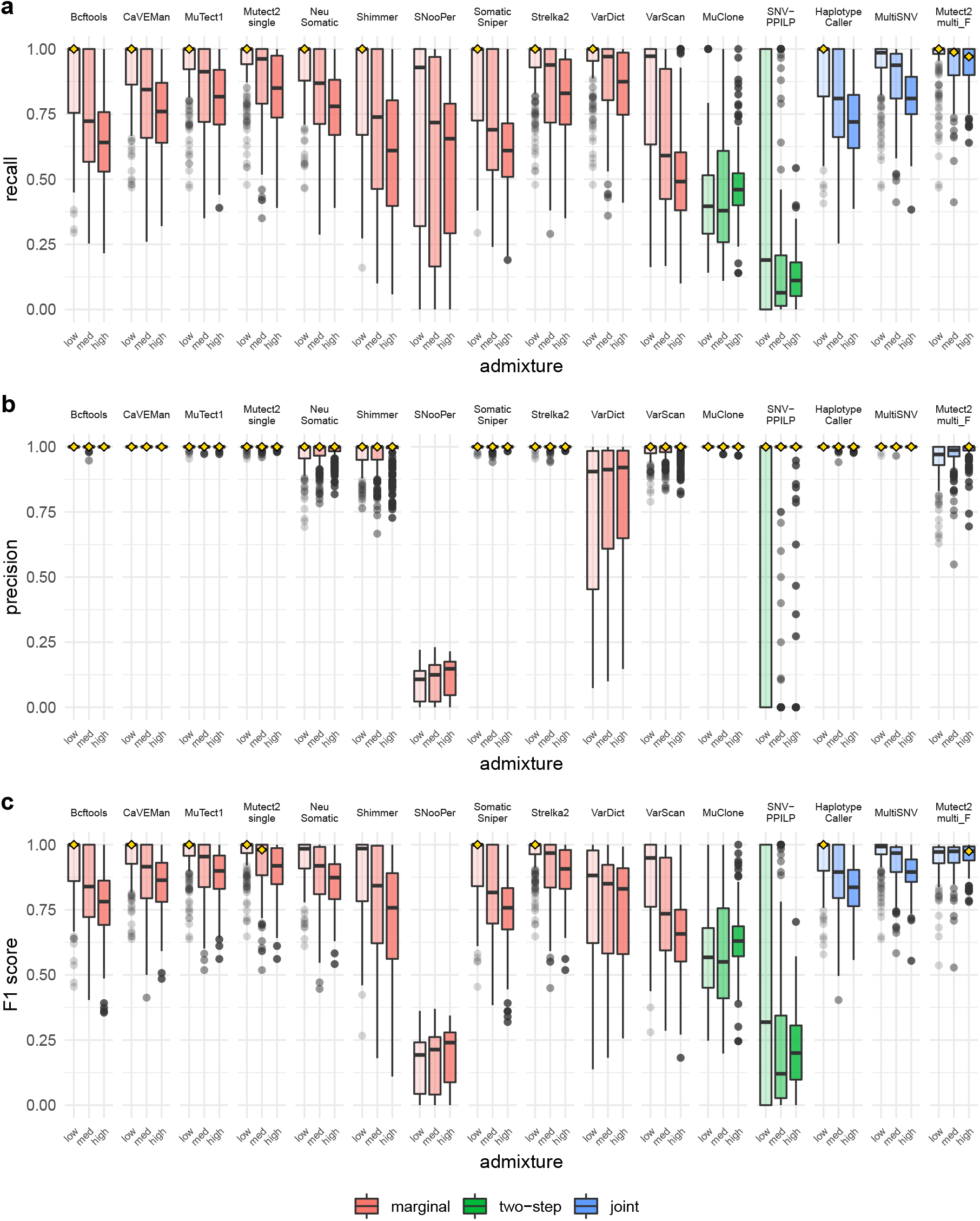
Performance of variant callers in the *de novo* simulation at different admixture levels. **a** recall. **b** precision. **c** F1 score. Boxplots as in Fig. 2. All metrics were calculated per tumor sample. The highest score in each category is highlighted with a yellow diamond.

### Effect of the variant allele frequency in the de novo simulation

The default filtering criteria for the different callers relate to thresholds above which variants were reported. In particular, the variant allele frequency (VAF) clearly affected recall (Fig. S1). True positive (TP) calls were most common for clonal variants (*V AF* = 0.5), but MultiSNV, the Mutect family, NeuSomatic, Strelka2 and VarDict were also quite good at low VAF. False negative (FN) variants were common for all methods at low VAF, but MuClone, SNooPer and SNV-PPILP showed also some FNs at higher VAF. False positive (FP) variants were absent in many callers, but were numerous mainly for clonal variants in SNooPer, and for low VAF in VarDict and Mutect2_multi_F. We analyzed in detail the origin of these FPs. SNooPer clearly detects many germline variants as somatic (Fig. S2). Most FPs in VarDict occur at high sequencing depth and with low alternative allele read counts, and seem to be the consequence of a lax filtering criterion, as by default two alternative reads already lead to a variant call. Mutect2_multi_F FPs were unrelated to germline variants.

### Overlap between variant call sets in the de novo simulation

Most callers detected similar sets of true variants, with the exception of SNV-PPILP, MuClone and SNooPer (Fig. S3). For the false negative calls there was less agreement. In terms of false positives, there was very little overlap among the callers. A hierarchical clustering of the callers based on the complete set of TP, FN and FP variant calls revealed similarities within the groups (MuTect1 / Mutect2_single / Mutect2_multi_F / Strelka2) and (CaVEMan / HaplotypeCaller / NeuSomatic), while SNooPer and SNV-PPILP were more dissimilar from all other methods (Fig. S4). To analyze which variant calls were caller-specific, and how they related to true variants, we compared all combinations of variant call sets. FP calls by SNooPer formed the largest set of variants (Fig. S5). The second largest set consisted of FN calls common to all methods, followed by VarDict FPs. The fourth largest set consisted of the TP calls of all methods apart from SNV-PPILP. The caller that detected the largest set of unique TPs was Mutect2_multi_F. Further set comparisons with the germline calls revealed that SNooPer FP calls largely consisted of germline variants, whereas VarDict FPs were almost exclusively somatic (Fig. S2). A comparison of the TP calls with true somatic variants revealed that the largest intersections contained most callers apart from SNV-PPILP, MuClone, and SNooPer (Fig. S6). Interestingly, Mutect2_multi_F and MultiSNV uniquely detected a large number of variants.

### Overall variant calling performance in the spike-in simulation

In general, the performance of the callers in the *spike*-*in* simulation was worse than in the *de novo* simulation. VarDict and the Mutect family showed the best recall, followed by NeuSomatic, HaplotypeCaller and SNV-PPLIP (Fig. 4). On the other hand, MuClone, Bcftools, CaVEMan, and particularly VarScan and SNooPer were clearly worse. The most precise caller was Shimmer (0.98), closely followed by the the Mutect family, and NeuSomatic. CaVEMan and SNooPer were highly imprecise. Mutect2_multi_F had the best F1 score, closely followed by Mutect2_single and MuTect1. Bcftools, CaVEMan, and particularly SNooPer, showed the worst F1 values. For MuTect1 and Mutect2_single, we observed a moderate increase in precision without a decrease in recall when using a panel of normals (PON) (Fig. S7).

**Figure 4:**
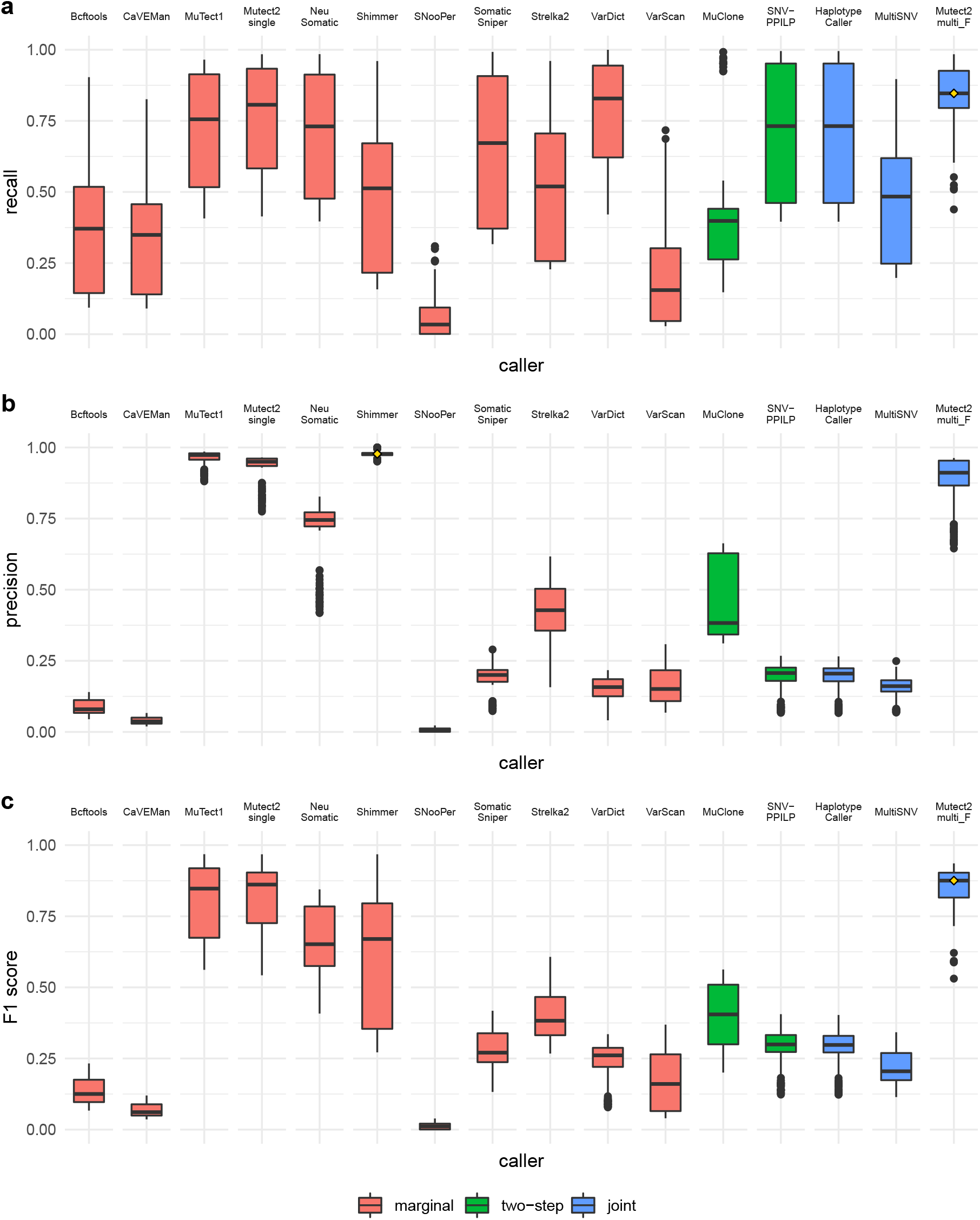
Performance of variant callers in the *spike*-*in* simulation. **a** recall. **b** precision. **c** F1 score. Boxplots as in Fig. 2. Metrics were calculated per tumor sample. The highest median score in each category is highlighted with a yellow diamond.

### Effect of clonal admixture in the spike-in simulation

Recall was consistently lower with higher admixture, except for MuClone and Mutect2_multi_F (Fig. 5). Precision increased with higher admixtures for MuTect1, Mutect2_single, NeuSomatic, Strelka2, VarDict, SNV-PPILP, HaplotypeCaller and Mutect2_multi_F, but decreased for Bcftools, CaVEMan, SNooPer and VarScan. The number of FPs was constant across admixture levels (Fig. S8). Remarkably, Shimmer was the best-performing caller at low admixture, while Mutect2_multi_F outperformed the others at medium and high admixture.

**Figure 5:**
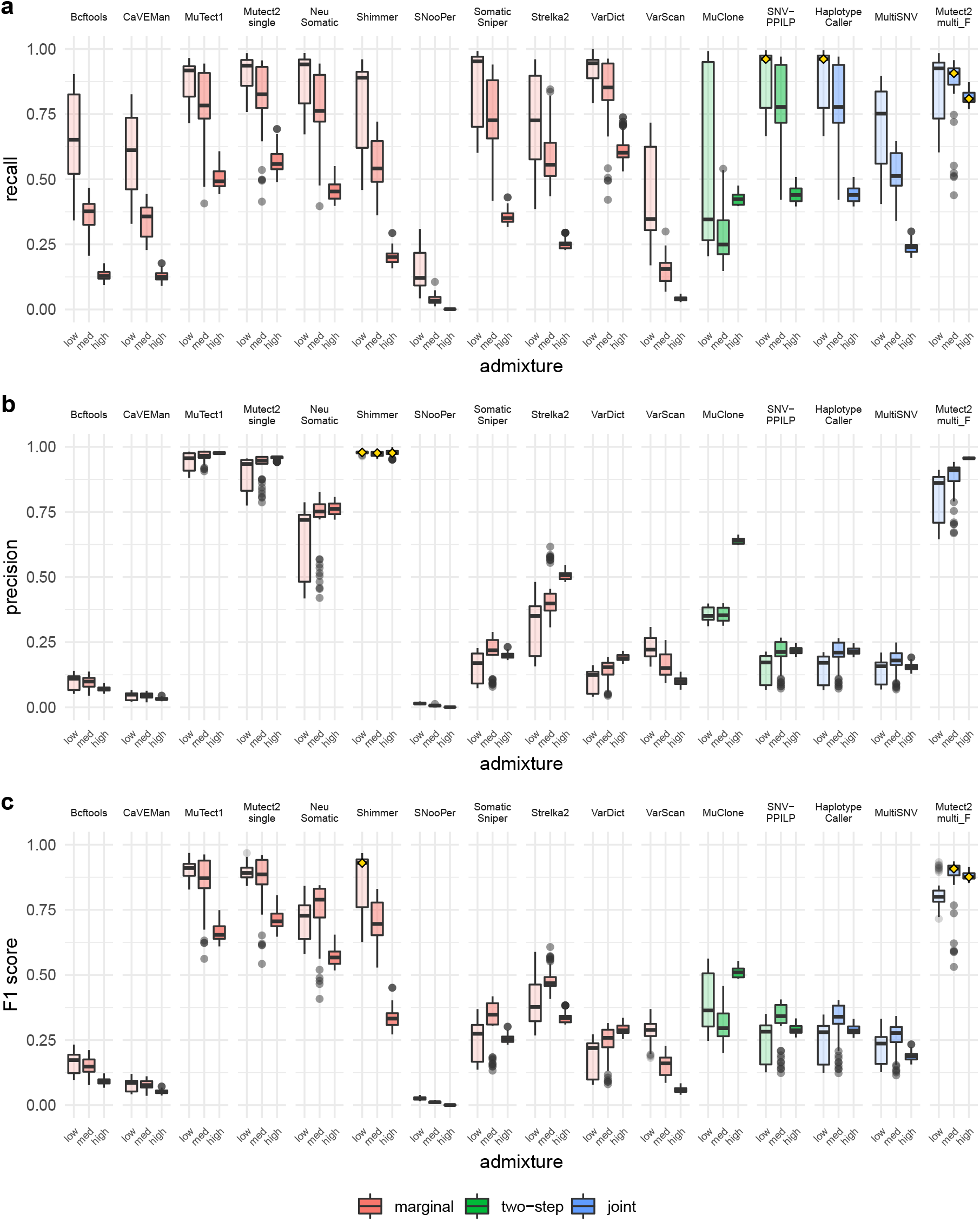
Performance of variant callers in the *spike*-*in* simulations at different admixture levels. **a** recall. **b** precision. **c** F1 score. Boxplots as in Fig. 2. Metrics were calculated per tumor sample. The highest median score in each category is highlighted with a yellow diamond.

### Effect of the variant allele frequency in the spike-in simulation

The VAF clearly influenced the TPs, which were most abundant for clonal variants. However, MuClone, the Mutect family, NeuSomatic, Strelka2 and VarDict seemed to have more TPs at relatively lower VAFs than other methods (Fig. S9). In most cases, FNs were more common for low-frequency variants, but to a different extent for the different callers. Most FPs arose at SNVs with relatively low frequencies, but Bcftools, HaplotypeCaller, SNooPer, SNV-PPILP, SomaticSniper and VarScan also reported FPs for high-frequency variants, a fraction of which were germline variants (22% for SNooPer, 25% for Bcftools and 26% for SomaticSniper) (Fig. S10a, Fig. S8). More than 20% of the FPs reported by HaplotypeCaller, SNV-PPILP, VarScan, and Shimmer were also called in the original 300x BAM file, but their allelic frequencies in the original 300x file and in the simulated 50x healthy sample were, in general, low (Fig. S10b,c). As expected, due to common sequencing errors, the FPs called in the same region in different replicates were significantly more similar than FPs called in different regions (Fig. S12).

### Overlap between variant call sets in the spike-in simulation)

When checking the overlap between call sets and true mutations (Fig. S13), FP calls dominated over all the sets. The largest set of TPs were those unique to Mutect2_multi_F (Fig. S14). The second largest TP set included all the callers but SNooPer. However, in the pairwise comparisons most methods but SNooPer called similar sets of TPs (Fig. S15a). FNs showed more disparity (Fig. S15b), with HaplotypeCaller and SNV-PPILP being the most similar ones. With some exceptions FPs were very different among callers (Fig. S15c, Fig. S8). A hierarchical clustering of the callers based on all the calls revealed similarities within the groups MuTect1/Mutect2_single/Mutect2_multi_F and HaplotypeCaller/SNV-PPILP, while SNooPer and CaVEMan were largely distinct (Fig. S16).

## Discussion

A general finding of our study was that MuTect1 and Mutect2_single consistently detected variants at high recall and precision, even if they have not been designed explicitly for the multiregional scenario. However, Mutect2_multi_F outperformed marginal callers in high-admixture scenarios, although it suffered from lower precision on the high-depth *de novo* tumors. MultiSNV worked quite well under ideal conditions (de novo simulations) on low-depth samples, in accordance with previous results[21], but performed significantly worse with the more noisy, empirical reads (spike-in simulations). While some of the tested callers achieved similar performance metrics in both the *de novo* and *spike*-*in* experiments, other methods performed considerably worse using empirical sequencing reads. When clonal admixture increased recall decreased, as expected due to the lower frequencies of subclonal variants. Precision was not expected to scale with admixture as FPs should be mostly independent of it, i.e., arising mainly from sequencing errors and germline variants.

Somewhat surprisingly, MuTect1 was slightly more precise than Mutect2_single on real sequencing data, despite not having realigned reads around indels, as recommended for the former[16]. Machine learning-based callers such as NeuSomatic and SNooPer are expected to depend on the quality of the training data used to derive the classification models. While NeuSomatic performed well under both simulation regimes, SNooPer showed many FPs, often coincident with germline variants (Fig. S1). SNV-PPILP had low recall rates; as it currently relies on the variant calls from HaplotypeCaller, its recall is limited by the output of HaplotypeCaller. VarDict detected many FPs at 300x, which can be attributed to its default settings, which means calling a variant if two reads support the alternative allele. MuClone had also very low recall, but when we fed it with the true clonal prevalences, its performance clearly improved (Fig. S17). Unfortunately this type of information is not usually available from real data and has to be estimated.

We used the default settings whenever possible. While the successive optimization of caller parameters typically yield significant improvements[18], the optimal parameterization is generally not obvious. In the *de novo* simulation, callers that need additional information (SNooPer needs a depth-specific classification model, SNV-PPILP assumes a perfect phylogeny and MuClone uses the PyClone clustering information), performed significantly worse than those that directly analyze the characteristics of the reads. Conversely, NeuSomatic, which uses a previously trained neural network for variant detection, worked quite well. An issue of general concern in benchmarking studies is whether results from simulated data generalize to empirical data sets[20]. Whenever ground truth information is required for a multitude of data sets, simulations are a useful tool. Simulated data by their very nature lack some of the features found in real data. In this study, for example, we focused on one type of mutation, SNVs, and did not introduce healthy cell DNA in the tumor samples. While those constraints limit the scope of the findings, they also represent a focused view of a controlled set of factors.

Here we strived to ensure the robustness of our findings by employing two different simulation approaches: 1) simulating sequencing reads in silico based on an artificial genome (de novo simulation), and 2) using an empirical sequencing data set and introducing somatic variants into them (spike-in simulation). *spike*-*in* simulations contain the noise inherent to real sequencing data, but they lack the absolute control achieved in the *de novo* simulations. Accurate somatic identification is fundamental for the study of somatic variation, including intratumoral heterogeneity, a fundamental feature of cancer. This study is the first one to provide a comprehensive comparison of different somatic variant callers in a multiregional scenario. It is also the first to compare recently released somatic variant callers like Mutect2, in particular its multisample mode, NeuSomatic, MuClone, MultiSNV and Strelka2. Importantly, we found that, given our simulation scenarios, ad hoc multiregional methods are not generally superior to some of the standard paired tumor-normal callers designed for simple samples. However, Mutect2 in multisample mode showed promising results, especially under difficult conditions - high admixture and low sequencing depth. If developers improve its genotyping of individual tumor samples, Mutect2 in multisample mode might easily become the leading method for M-seq tumor data; as of the time of this study, users are advised to apply additional filters to avoid false positive calls when using this method.

Taking the criterion of robustness under different conditions into account, Mutect2_multi_F could be the most efficient strategy so far for the analysis of multiregional sequencing data coming from high-admixture tumors, for example liquid tumors, particularly when high recall is desired. On the contrary, for low-admixture tumors like adenomas and some colorectal carcinomas[22, 23], single-sample callers like Mutect2_single or Shimmer might also perform well.

## Methods

We employed two different simulation approaches to generate M-seq data; 1) a completely controlled *de novo* simulation, sampling synthetic reads from randomly generated genomes, followed by introduction of germline and somatic single-nucleotide variants (sSNVs), and 2) a simulation based on real sequencing data split into multiple samples, followed by the *spike*-*in* of simulated sSNVs into real reads.

### De novo simulation of synthetic reads

For each tumor data set, we constructed a tree (Fig. 1a) with six clones by random step-wise addition. We fixed the number of regions to six (one healthy plus five tumoral), and distributed the clones across them according to three predefined scenarios: 1) low-admixture (few clones per sample, expected little clonal overlap); 2) moderate-admixture (moderate number clones per sample, expected moderate clonal overlap); and 3) high-admixture (high number clones per sample, expected high clonal overlap) (Fig. 1b, Fig. S18). We picked the first clone for each sample randomly and then selected the co-occurring clones based on tree distance. Then we sampled the prevalence of each clone at each sample from a flat Dirichlet prior. All samples were simulated to be pure, that is, we assumed no normal cell contamination. More details about this simulation in Supplementary Information.

We generated 120 tumor replicates. For each replicate, we simulated a haploid genomic reference sequence of 3 Mb by randomly sampling bases according to the nucleotide frequencies of the human genome (hg19). To produce a healthy, diploid human genome from the haploid reference, we generated 3000 germline single nucleotide polymorphisms (SNPs) under a transition/transversion ratio of 1.7, and assigned them to the maternal or paternal genome (heterozygous), or to both (homozygous) with a probability of 0.2 (Fig. 1f). We then generated a set of 100 sSNVs according to the trinucleotide mutational signature 1 from COSMIC^1^. We selected signature 1 because it is common in all cancer types. Subsequently, we assigned each sSNV to either the paternal or maternal allele and placed it on a random branch of the clone tree (Fig. 1c).

We simulated PE100 Illumina reads from the reference genome for each regional sample to an average depth of 30, 50, 100 or 300x using ART[24] (Fig. 1e). Finally, we assigned each read pair to a clone (either the maternal or the paternal allele) according to the prevalence matrix. Depending on the presence of SNPs and sSNVs in the chosen clone and allele, we introduced the simulated sSNVs into the reads (Fig. 1d). For each combination of admixture scenario and sequencing depth a set of ten replicates was generated, amounting to a total of 120 M-seq tumor data sets.

### Spike-in simulation of variants in real reads

We downloaded the 300x WGS (TruSeq DNA PCR-Free and Illumina PE250) FASTQ files corresponding to twelve sequencing libraries for sample NA24631 from the Genome in a Bottle (GIAB) FTP server^2^ (Fig. 1m,k). These data come from a large homogenized culture of a B lymphoblastoid cell line, derived from the healthy blood of a 33-year old chinese man. We trimmed the sequencing adapters using Cutadapt[25] (v.1.15) and filtered out reads shorter than 70 bp. We aligned the reads to the human reference genome hs37d5 using BWA-MEM[26] (v.0.7.17) and sorted them using Picard^3^ (v.2.2.1) SortSam. We utilized Picard MarkDuplicates to remove duplicates and to simultaneously merge the files from the different libraries. To make the simulations computationally feasible, we used samtools[26] view (v.1.9) to keep only reads mapped to chromosome 21. We chose this chromosome because it is the smallest autosome, its GC content is representative of the whole genome and it has a lower number of troublesome sequences for alignment^4^.

We recreated a multiregional sampling scheme by splitting the original bam file into six different regions (Fig. S19). In order to introduce mutations always in the same phase regarding the heterozygous germline variants (i.e., single nucleotide polymorphisms or SNPs) already present in this genome, we first pseudo-phased the original bam file with phase-tools^5^ (v.1.1.3). Phase-tools uses heterozygous SNPs to separate the reads into two different bam files (maternal and paternal). We worked with the two files independently for the remaining steps of the simulation. Reads with an ambiguous phase were randomly assigned to one of the maternal/paternal BAM files. Afterwards, we randomly split the reads into six equally-sized files: one healthy sample and five tumor regions (Fig. 1l). Each read was assigned to exactly one region. The high depth of the original data set allowed us to produce 50x regional samples with unique reads.

In total, we generated 30 different M-seq data sets combining ten different mutation matrices and three clonal admixture scenarios (Fig. S20). For every data set, we introduced 1000 SNVs distributed among seven clones (Fig. 1i). In all cases, the tumoral clones shared a common evolutionary history defined by a single, fixed tree topology with populated internal nodes and equal branches (Fig. 1g), so the SNVs were distributed among the seven clones accordingly. The mutation matrices reflect the physical positions of the mutations. We chose the alternative alleles and the positions to mutate according to COSMIC mutational signature 5^6^. We selected signature 5 because it is very common in all cancer types. To introduce mutations according to this signature (Fig. S21), we first obtained all the trinucleotides in the healthy NA24631 consensus maternal and paternal haplotypes. Then, we selected a type of trinucleotide change according to its probability in signature 5, and added it to the mutation matrix. As the nucleotides overlap, when the central position of a trinucleotide changes, not only such trinucleotide is affected, but also the previous and the next one. This means that all three trinucleotide now belong to a different class, and will have a new change probability. We chose to remove them from the list of trinucleotides available to mutate for computational convenience. In practice, this means that we impeded mutations occurring on overlapping trinucleotides, something very unlikely in any case for 1000 sSNVs distributed along chromosome 21 (48.13 Mb).

The three admixture scenarios corresponded to low, medium or high clonal admixture. We defined them using three arbitrary prevalence matrices that reflected the clonal composition of each tumor region (Fig. 1h). To simulate each admixture scenario, we randomly split the BAM file for each region into clones according the proportions in the corresponding prevalence matrix. For each replicate, we introduced the SNVs of the mutation matrix to the corresponding clone using BamSurgeon[18] (v.1.1) (Fig. 1j). Since each file represents one allele of one clone, we introduced the SNVs at *V AF* = 1.0. Finally, we merged back both alleles of all the clones belonging to each region and modified the read groups accordingly.

### Variant calling

We considered published variant calling methods that we grouped in three arbitrary categories regarding how we used them with M-seq data: marginal, joint, and two-step calling (Table 1). Under the marginal calling strategy we included popular somatic variant callers that are typically used for identifying sSNVs in single tumor-normal paired samples. Here we used these tools in a multiregional setting simply applying them independently for each of the regions simulated. Under the two-step calling strategy we included MuClone[12] and SNV-PPILP[13]. MuClone incorporates prior knowledge about the clonal composition of the tumor to classify a given set of candidate variants; SNV-PPILP assumes all samples to be monophyletic and applies the perfect phylogeny assumption to filter a given set of variants. Under the joint calling strategy we considered MultiSNV[14], that employs a joint statistical model across samples, Mutect2 multi, that aggregates the reads from all tumor samples, and HaplotypeCaller[15] that focuses on germline variants in population-wide data sets, but we think it may be applicable to somatic cell populations within individuals, hence we included it in the comparison.

**Table 1:** Multiregional somatic variant calling strategies included in this study.

| Tool | Citation | M-seq calling strategy |
| --- | --- | --- |
| Bcftools [27] | Li et al., 2011 | marginal |
| CaVEMan [28] | Jones et al., 2016 | marginal |
| MuTect1 [16] | Cibulskis et al., 2013 | marginal |
| Mutect2 [16] | Cibulskis et al., 2013 | marginal / joint |
| NeuSomatic [29] | Sahraeian et al., 2019 | marginal |
| Shimmer [30] | Hansen et al., 2013 | marginal |
| SNooPer [31] | Spinella et al., 2016 | marginal |
| SomaticSniper [32] | Larson et al., 2012 | marginal |
| Strelka2 [33] | Kim et al., 2018 | marginal |
| VarDict [34] | Lai et al., 2016 | marginal |
| VarScan2 [35] | Koboldt et al., 2012 | marginal |
| HaplotypeCaller [15] | Poplin et al., 2018 | joint |
| MultiSNV [14] | Josephidou et al., 2015 | joint |
| MuClone [12] | Dorri et al., 2017 | two-step |
| SNV-PPILP [13] | van Rens et al., 2015 | two-step |

### Variant calling parameters

MuTect1, Mutect2_single, Mutect2 multi, SomaticSniper, Strelka2, Shimmer, MultiSNV, VarScan and VarDict take tumor and healthy BAM files plus the reference genome as input. SNooPer and VarScan require pileup files, so we converted the BAM files to pileup using samtools mpileup 1.8 (the current standard bcftools mpileup is not supported by these callers). Strelka2 needed a simple configuration step to set up the input files and working directory. SNV-PPILP works with the HaplotypeCaller output, so we ran GATK-HaplotypeCaller for each sample in GVCF mode, aggregated the multi-sample information with GenomicsDBImport, performed joint genotyping with GenotypeGVCFs, and filtered out the variants that were not homozygous for the reference allele in the healthy sample. MuClone requires variant calls from another caller, so we supplied it with the unfiltered calls from Mutect2_single (marginal calling mode) and we ran it setting the parameter *phi_threshold* to 0.01. MuClone also needs the clustering information of PyClone, so we ran PyClone using the filtered biallelic calls from Mutect2_single (we used the unfiltered calls as MuClone input to avoid biasing the calling, and the filtered ones as PyClone input to obtain higher-accuracy clusters). The median depth for MultiSNV was set to the one simulated in the *de novo* simulation, and to 49x (the median depth calculated with samtools depth) in the *spike*-*in* simulation. In the latter case, we limited the analysis to chromosome 21 for the callers that support region-specific execution. We used the default values for the remaining parameters.

Working with real human sequencing data in the *spike*-*in* simulations allows the callers to take advantage of external information. For MuTect1, Mutect2_single and Mutect2 multi we used a panel of normals (PON). Ideally, PONs are built from a set of healthy samples obtained with the same sequencing workflow as the tumor samples to analyze. All the variants that appear in more than one sample of the PON are assumed to be either common SNPs or recurrent sequencing errors, so they will be filtered out during variant calling. In this case, to build the PON we downloaded 66 healthy samples sequenced with TruSeq and Illumina Hiseq from The 100-Person Wellness Project (HPWP[36] from dbGAP. We ran MuTect1 and Mutect2_single both with and without the PON to assess its effect. We also provided the Mutects with known sites to filter out germline variants that are common in the population. Specifically, we used dbSNP 138 and Cosmic v83 databases for MuTect1 and gnomad for Mutect2_single and Mutect2 multi.

### Variant filtering

VarDict needs a specific filtering of the identified variants due to its “calling everything” philosophy (https://github.com/AstraZeneca-NGS/VarDict), so we ran the testsomatic.R script included in VarDict. In the case of Mutect2 (tumor-normal and multisample modes), GATK provides its own filtering tool called FilterMutectCalls, which we executed with the default parameters. Additionally, for Mutect2_multi we filtered out all variants without reads for the alternative allele, because at the time of this study, and according to the authors the tool reported variants tumor-wide, but not proper genotypes for each sample^7^. Thus, herein we refer to the results of that approach as “Mutect2_multi_F”. MuTect1, SNooPer, Strelka2, VarDict, MultiSNV, SNV-PPILP, NeuSomatic and VarScan perform their own internal filtering, so in this case we selected the variants tagged as PASS. Besides, SomaticSniper VarDict, Multi-SNV, and VarScan report not only somatic but also germline variants or LOH events. We used the field “somatic status” to select only the somatic calls (StrongSomatic and LikelySomatic tags in the case of VarDict). Finally, we used GATK SelectVariants to remove indels by selecting SNPs and MNPs (multi-nucleotide polymorphisms) types.

### Performance metrics

We measured the variant calling performance in terms of recall and precision, which are derived from the tally of true positives (TP), false positives (FP) and false negatives (FN). Recall is the proportion of the true variants that is captured (*TP*/(*TP* +*FN*)), while precision is the proportion of the captured variants that is true (*TP*/(*TP* + *FP*)). We defined as TP any variant called in a regional sample where it is actually present. Conversely, an FP will be any variant called in a regional sample where it is not present. In the same way, an FN will be any variant that is not called in a regional sample where it is present. Note that FN calls measure the effect of sensitivity of the respective variant caller as well as the sampling bias at low sequencing coverage. Herein, “presence” of a variant is determined by the prevalence matrix, i.e., if a clone occurs in a sample, the mutations it carries are considered to be present in that sample. Finally, we also computed an overall measure that combines recall and precision, the F1 score (*F*1 = 2 ∗ *recall* ∗ *precision*/(*recall* + *precision*)).

## Acknowledgements

This work was supported by the European Research Council (ERC-617457- PHYLOCANCER awarded to DP) and by the Spanish Ministry of Economy and Competitiveness - MINECO (BFU2015-63774-P awarded to DP). DP receives further support from Xunta de Galicia. LT is supported by a Ph.D. fellowship from Xunta de Galicia (ED481A-2018/303). TP is now supported by a Ph.D. fellowship from the Spanish Government (FPU15/03709) and previously by a Ph.D. fellowship from Xunta de Galicia (ED481A-2015/083). We thank Andrés Pérez-Figueroa for valuable advice as to visualization of the results, João Alves for support in the interpretation of variant caller output, Fabian Crespo for discussion of performance metrics, Nathan D. Price for providing the healthy samples to build the PON and the Supercomputing Center of Galicia (CESGA) for computing services.

## Author contributions

DP supervised the whole study. HD and DP designed the *de novo* simulations and HD implemented them. LT, TP and DP designed the *spike*-*in* simulations, which were carried out by LT and TP; HD and LT ran the variant calling methods and analyzed the results with help from TP. HD, LT and DP wrote the manuscript. All authors read and approved the final manuscript.

## Data Availability

All data sets simulated in this study can be reproduced/obtained using the scripts available in the GitHub repository https://github.com/hdetering/mseq-vc. The FASTQ files for sample NA24631 can be obtained from the Genome in a Bottle (GIAB) FTP server^8^.

## Code Availability

The code for simulation and analysis of data supporting the conclusions in the article are available in the GitHub repository https://github.com/hdetering/mseq-vc.

## Footnotes

1 https://cancer.sanger.ac.uk/cosmic/signatures_v2

2 https://ftp-trace.ncbi.nlm.nih.gov/giab/ftp/data/

3 http://broadinstitute.github.io/picard/

4 https://genome.ucsc.edu/

5 https://github.com/mateidavid/phase-tools

6 https://cancer.sanger.ac.uk/cosmic/signatures_v2

7 https://gatkforums.broadinstitute.org/gatk/discussion/comment/56475/#Comment_56475

8 https://ftp-trace.ncbi.nlm.nih.gov/giab/ftp/data/

