## Supplementary Material for "Accuracy of somatic variant detection in multiregional tumor sequencing data"

Harald Detering[1,2,3,e], Laura Tomás[1,2,3,e], Tamara Prieto[1,2,3], and David Posada[1,2,3,*]

[1]*Department of Biochemistry, Genetics, and Immunology, University of Vigo, Spain.*
[2]*Biomedical Research Center (CINBIO), University of Vigo, Spain.*
[3]*Galicia Sur Health Research Institute, Vigo, Spain.*
[e]*These authors contributed equally*
[*]**

### 1 Supplementary Methods

#### 1.1 De novo simulation

The following subsections correspond to the panels in Fig. 1 in the main text. All steps were carried out for each of 120 replicates of the de novo simulation. Workflow steps are described in order of execution, rather than by lexicographical order of panels in Fig. 1 in the main text.

##### 1.1.1 Simulating a healthy genome

In order to generate a healthy genome, a reference genome sequence of 3 Mb size was simulated for each tumor by generating a random sequence of nucleotides according to the ACGT frequencies in the human reference genome hg19. Then, we added 3000 germline mutations (i.e., 1 SNP every 1000 sites) to this sequence assuming a transition/transversion ratio of 1.7. The genomic location of the chosen mutation was selected at random according to the nucleotide frequencies. We enforced the infinite sites assumption, i.e., mutations were always assigned to a location not mutated previously. Each mutation was assigned to either the maternal or paternal chromosome, or to both in the case of homozygous mutations. The heterozygous/homozygous status of each mutation was picked at random with a 0.2 probability of being homozygous (a similar value to that observed in the healthy sample described below for the spike-in simulations).

#### 1.1.2 Generation of clone tree for a given number of clones

For a fixed number of clones (six in this case), we constructed a tree, with populated internal nodes and possible polytomies, using Algorithm 1 below (code available at:
https://github.com/hdetering/mseq-vc/blob/master/scripts/de-novo.sim-tree-prev.ipynb
https://github.com/hdetering/mseq-vc/blob/master/scripts/de-novo.sim-tree-prev.html.
Each tree was then used to generate prevalence matrices and assign somatic mutations in later steps.

---

**Algorithm 1:** Simulation of clone trees

---

**input :** number of clones $n$
**output:** Nodes $N$, Parent relationship *parent*

**init:** *Healthy node* $n_h$
*Set of tree nodes* $N := \{n_h\}$
*Parent relationship parent* $:= \{(n_h, NULL)\}$
*Set of available parent nodes* $A := \{\}$

Add new node $c_1$ to $N$. Assign $parent(c_1) = n_h$. Add $c_s1$ to $A$.
**for** $i \in \{2, \ldots, n\}$ **do**
  Add new node $c_i$ to $N$
  Pick random node $p$ from $A$; assign $parent(c_i) = p$
  Add $c_i$ to $A$
**end**

---

#### 1.1.3 Generating clonal mixtures for each sample

In order to allocate the clones in the different regions, we defined three different admixture scenarios (Figure S13):

- Low admixture: higher probability for fewer clones per sample
- Medium admixture: identical probability for any number of clones (1-6) per sample
- High admixture: higher probability for many clones per sample

We assumed that more closely related clones are more likely to occur in the same sample. Hence, we used the distance between clones in the clone tree (i.e., the sum of branch lengths in the path connecting both nodes) as a measure of relatedness. Clonal locations and prevalences were then assigned using Algorithm 2 below; code available at:
https://github.com/hdetering/mseq-vc/blob/master/scripts/de-novo.sim-tree-prev.ipynb
https://github.com/hdetering/mseq-vc/blob/master/scripts/de-novo.sim-tree-prev.html

#### 1.1.4 Generating somatic mutations

For each tumor, we generated a total of 100 somatic single nucleotide polymorphisms (sSNV). The type and the location of each mutation was picked at random using of the trinucleotide Signature 1 available at COSMIC https://cancer.sanger.ac.uk/cosmic/signatures_v2. Each mutation was assigned to a random branch in the clone tree.

**Algorithm 2:** Generating clonal multiregional prevalences

**input :** Set of clones $C$
Relatedness matrix $R$ ($C \times C$, values $exp(-d)$ for tree distance $d$)
Prior probabilities $p$ for the number of clones per sample

**output:** Prevalence matrix $F(S \times C)$, assigning a cellular frequency to every clone in every sample

**for** *regional sample* $s \in \{R1, \ldots, R5\}$ **do**
  **init:** *Set* $A = C$ *of available clones in sample* $s$
  *Set* $I = \{\}$ *of included clones in sample* $s$

  Select random number of clones $n$, using $p$ as weights
  Select random clone $c_1$, add $c_1$ to $I$, remove $c_1$ from $A$
  **for** $i \in \{2, \ldots, n\}$ **do**
    Select random clone $c_i$ from $A$ with weights $R(c_1)$
    Add $c_i$ to $I$, remove $c_i$ from $A$
    Pick random prevalence values $(f_1, \ldots, f_n)$ using a flat Dirichlet prior
    Assign prevalence 0 to all clones in $C \setminus I$
  **end**
**end**

#### 1.1.5 Simulating sequencing reads

For each sample and clone, we simulated sequencing reads from the reference genome of each tumor using the sequencing simulator ART[1] v2.5.8. The sequencing coverage was chosen in proportion to each clone's cellular frequency as specified in the prevalence matrix. All other parameters were identical for all runs, namely: `--len 100`, `--paired`, `--mflen 500`, `--sdev 20`, `--seqSys HS25`. That is, we simulated paired-end reads of length 100bp with a fragment size of $500 \pm 20bp$ based on the error profile of an Illumina HiSeq 2500 machine.

#### 1.1.6 Adding mutations to sequencing reads

After reads were generated for each clone, they were merged for each regional sample into a bulk tumor sequencing dataset. In the process of merging, each read pair was assigned to the maternal or paternal allele. Further, germline mutations and somatic variants were added to the reads if they affected the allele and the clone that the read pair originated from. Finally, reads for each sample were aligned against the tumor reference genome using BWA-MEM[2]. As the end result, each tumor was represented by five BAM files which include germline mutations and somatic single-nucleotide variants.

### 2 Supplementary figures

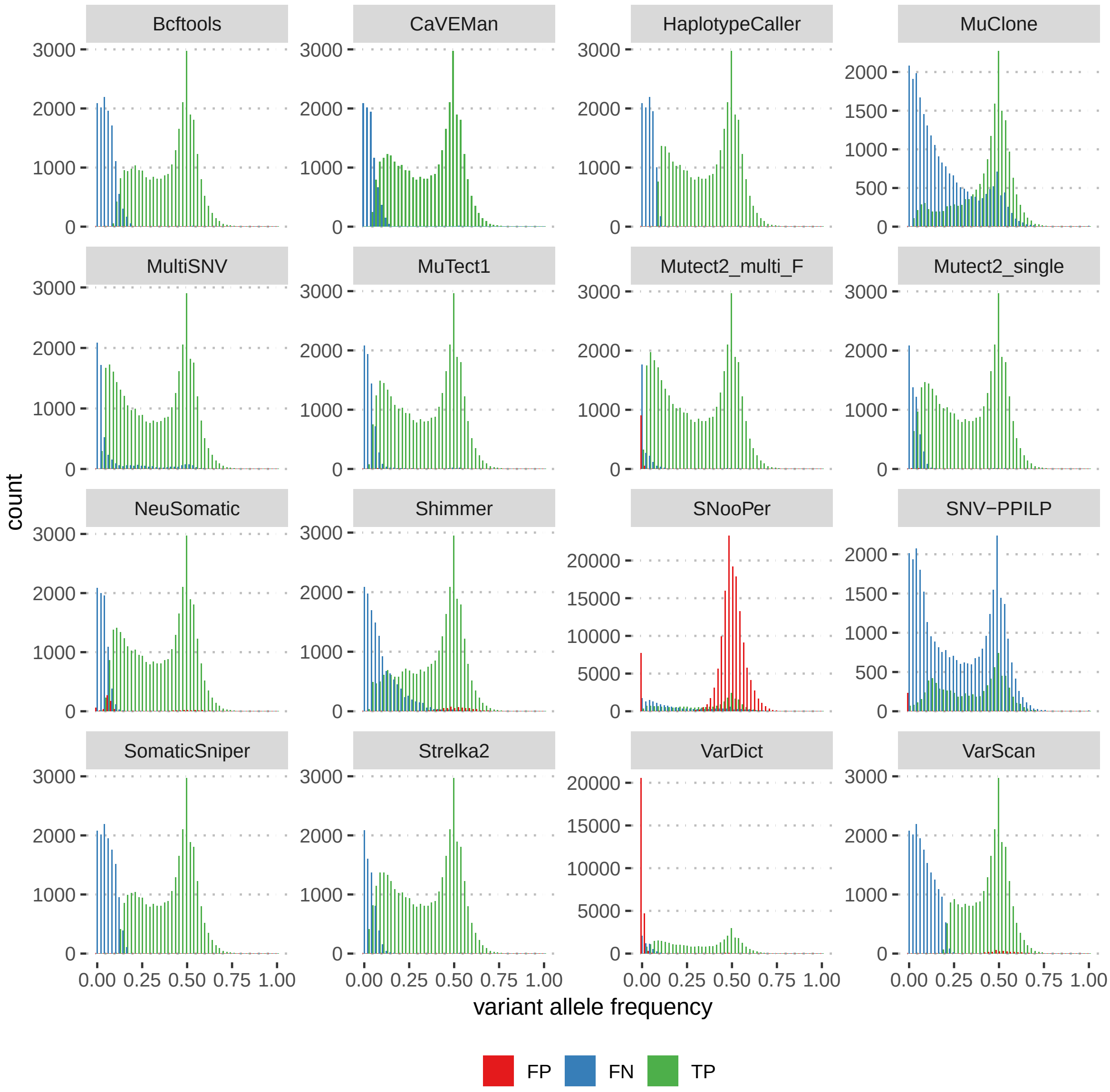


Figure S1: Variant allele frequencies in the *de novo* simulation for true positive (TP), false positive (FP) and false negative (FN) calls. Y axes scaled individually for each subgraph.

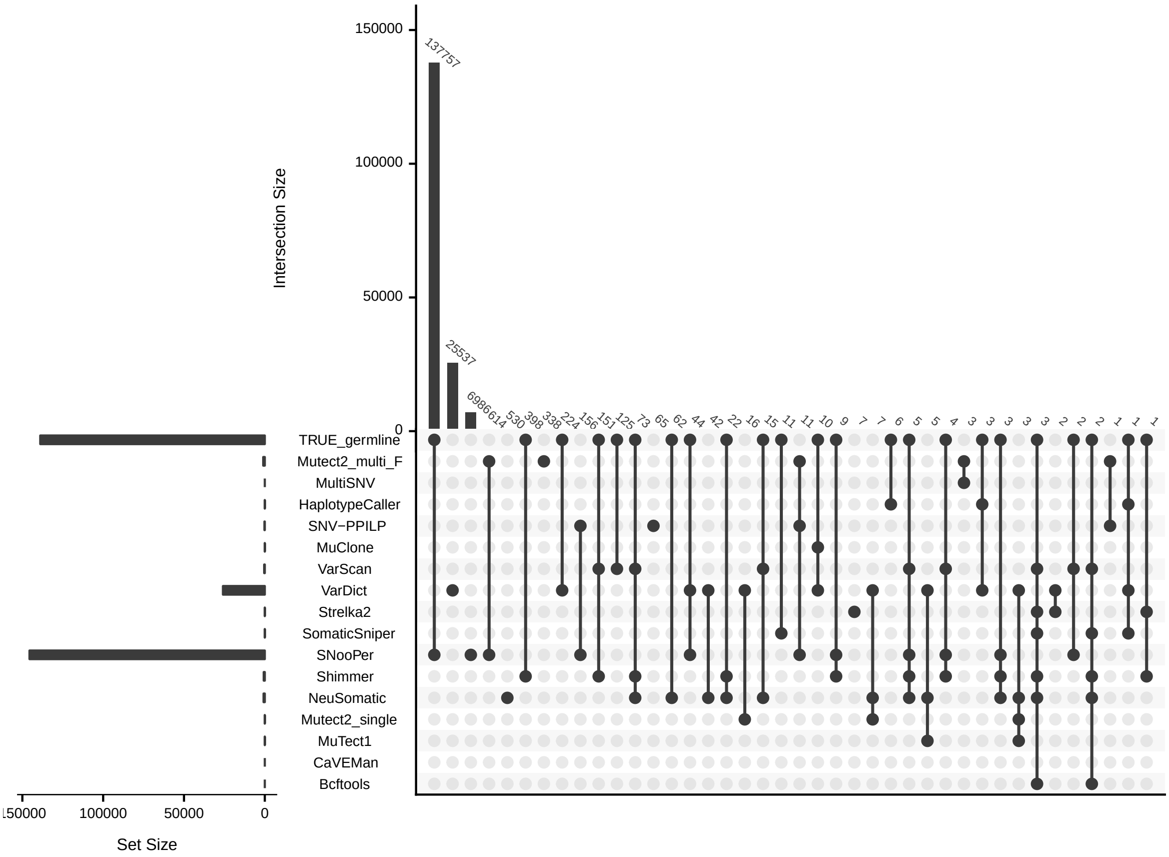


Figure S2: Intersection between false positive (FP) calls and germline variants in the *de novo* simulations. Every row represents a single set of variants, and the horizontal bars on the left indicate the size of these sets. Every dot column represents a given intersection between sets, and the thick vertical bars on top show the size of these intersections. The black dots (connected by black lines) show the sets involved in each intersection. Only the largest 40 intersections are shown. The TRUE_germline set contains the simulated SNPs, while the rest contains the set of variants called by each method.

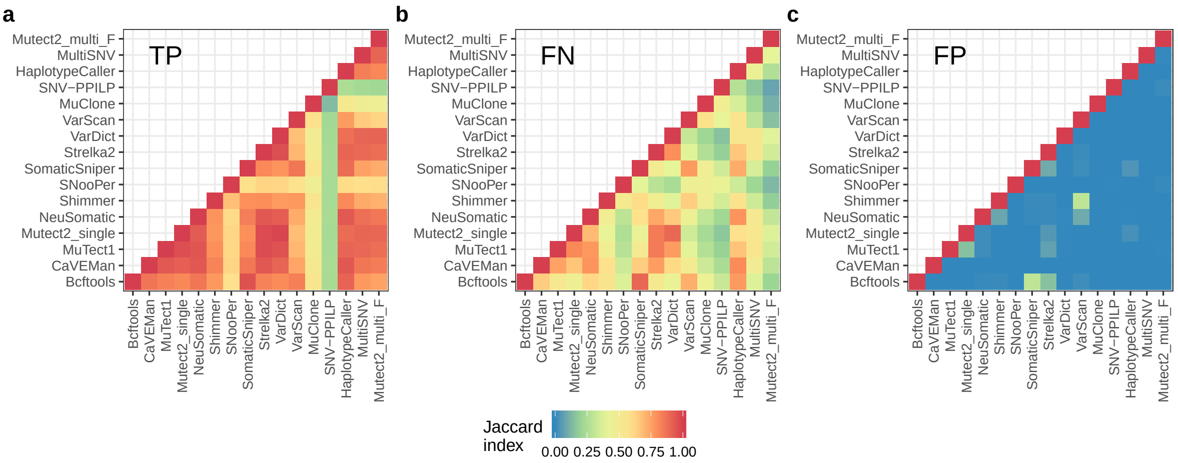


Figure S3: Jaccard distance between variant call sets in the *de novo* simulations. **a** True positive (TP). **b** False negatives (FN). **c** False positives (FP).

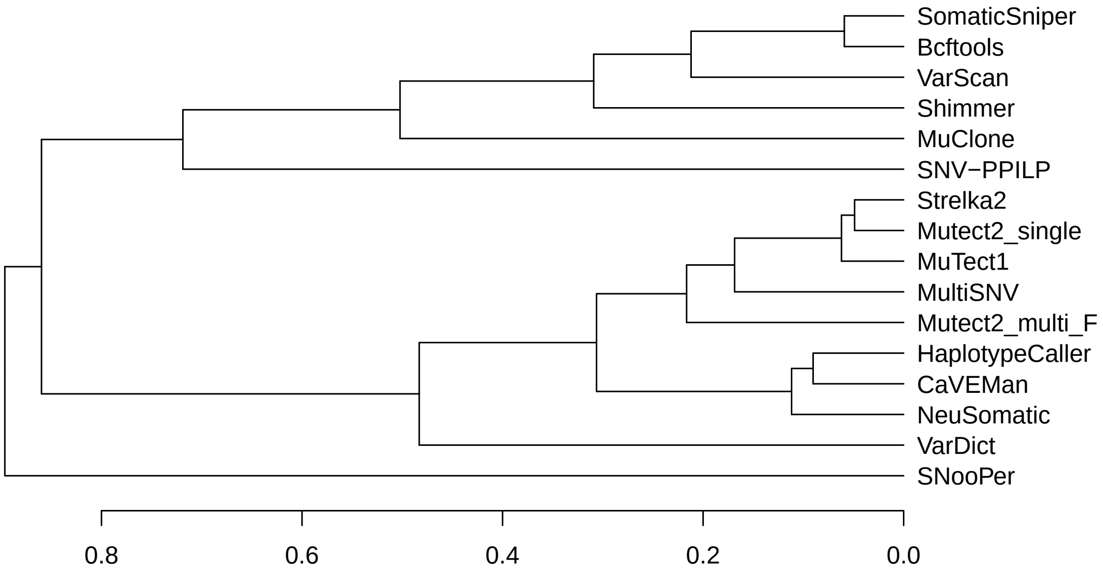


Figure S4: Hierarchical clustering of callers in the *de novo* simulations. The clustering was carried out according to the Jaccard distance between the variant call sets for the different callers.

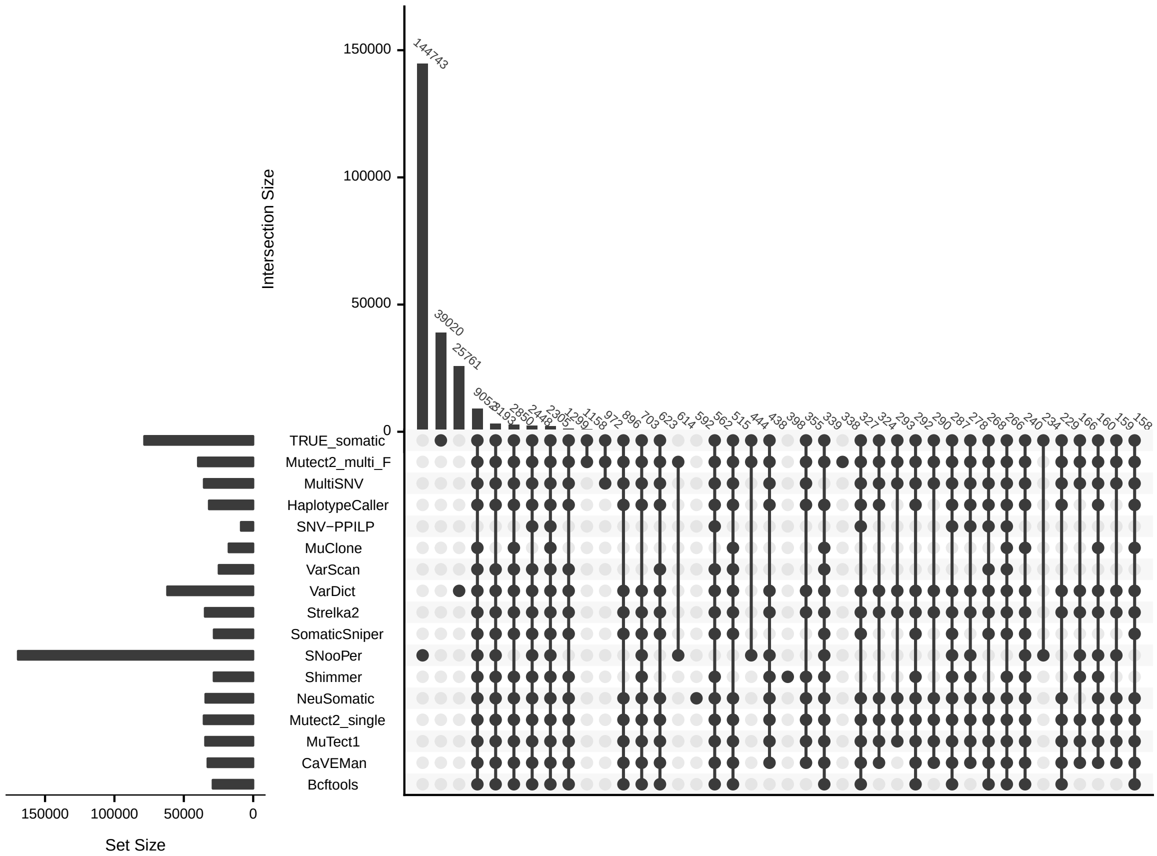


Figure S5: Overlap between variant calls and true somatic variants in the *de novo* simulations. TRUE_somatic is the set of simulated somatic mutations. See Fig. S2 for a detailed explanation of the figure components.

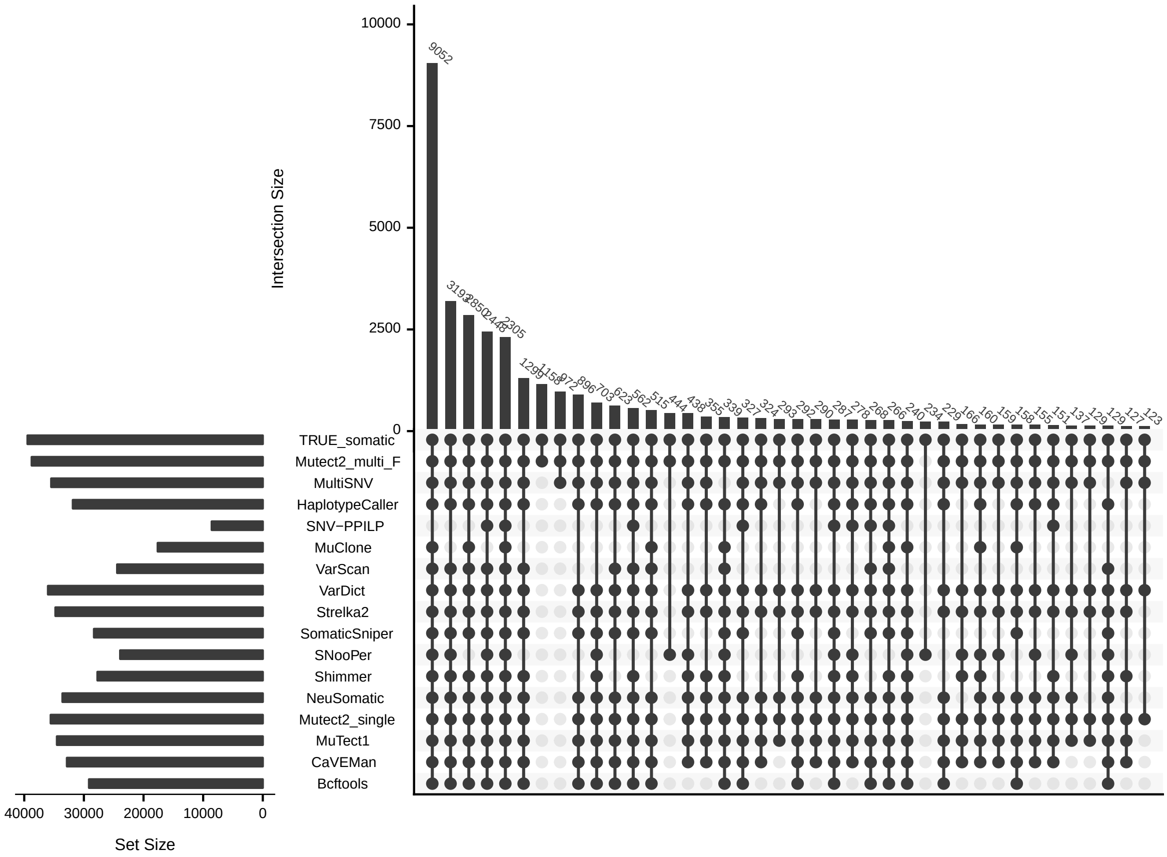


Figure S6: Intersection between true positive (TP) calls and true somatic variants in the *de novo* simulations. TRUE_somatic is the set of simulated somatic mutations. Only intersections involving TRUE_somatic are shown, so only the intersections between TP sets are observed. See Fig. S2 for a detailed explanation of the figure components.

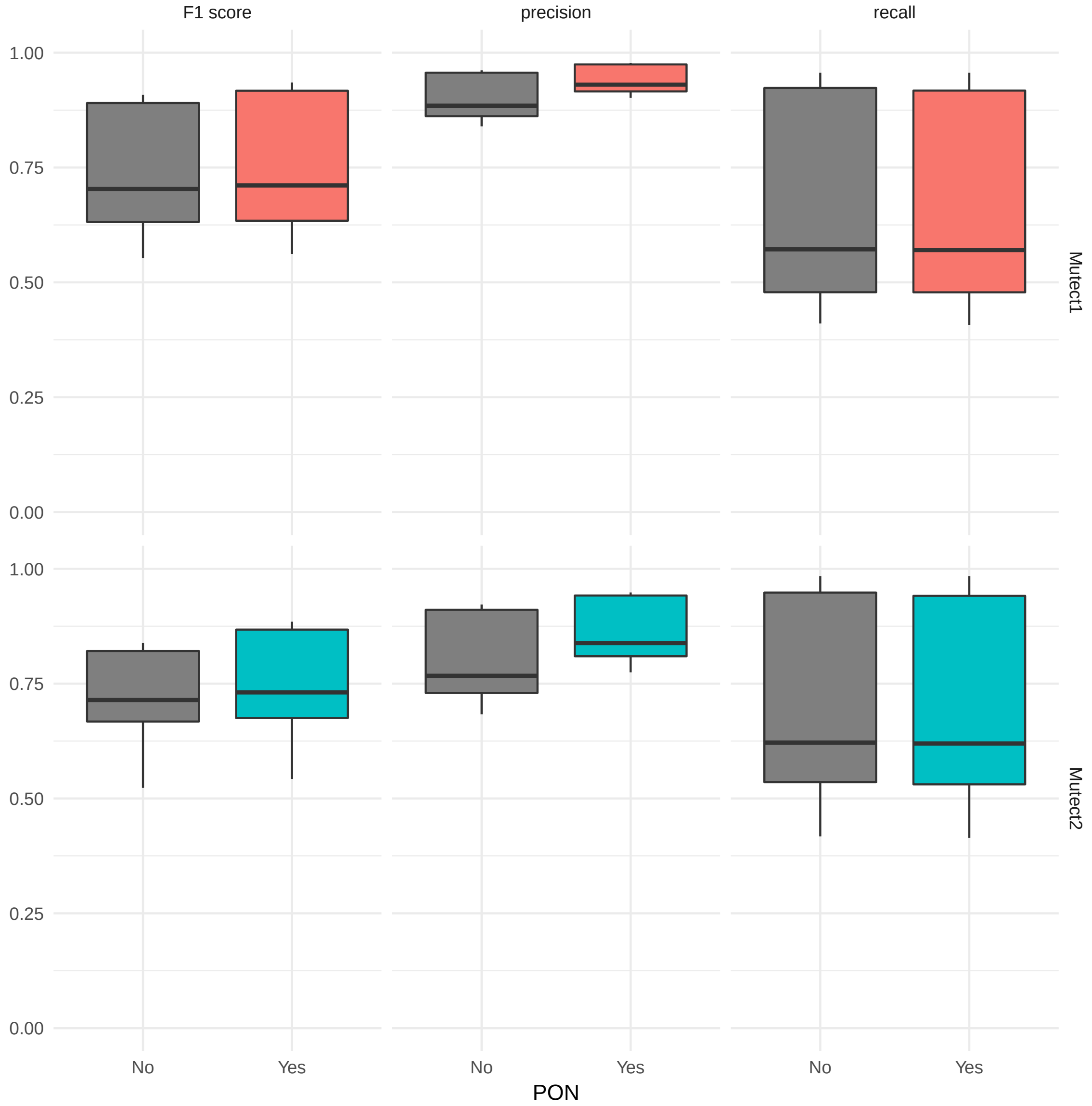


Figure S7: Effect of the panel of normals on the performance of MuTect1 and Mutect2_single. Recall, precision and F1 score are shown for MuTect1 and Mutect2_single with and without a set of recurrent sequencing and library errors and common SNPs previously identified in a set of 66 healthy samples (panel of normals or PON).

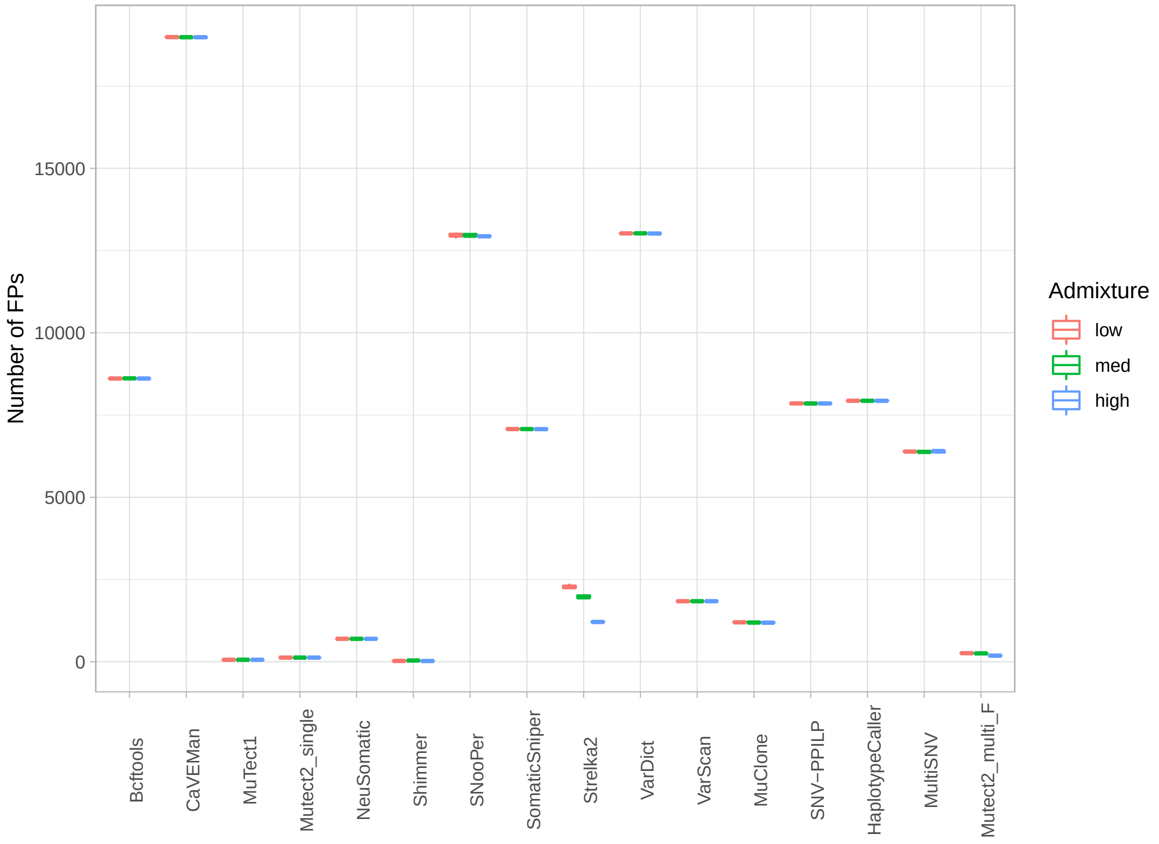


Figure S8: Effect of admixture in the number of false positives in *spike-in* simulation. Each boxplot represents the number of FPs in ten replicates with low, medium or high admixture for each caller.

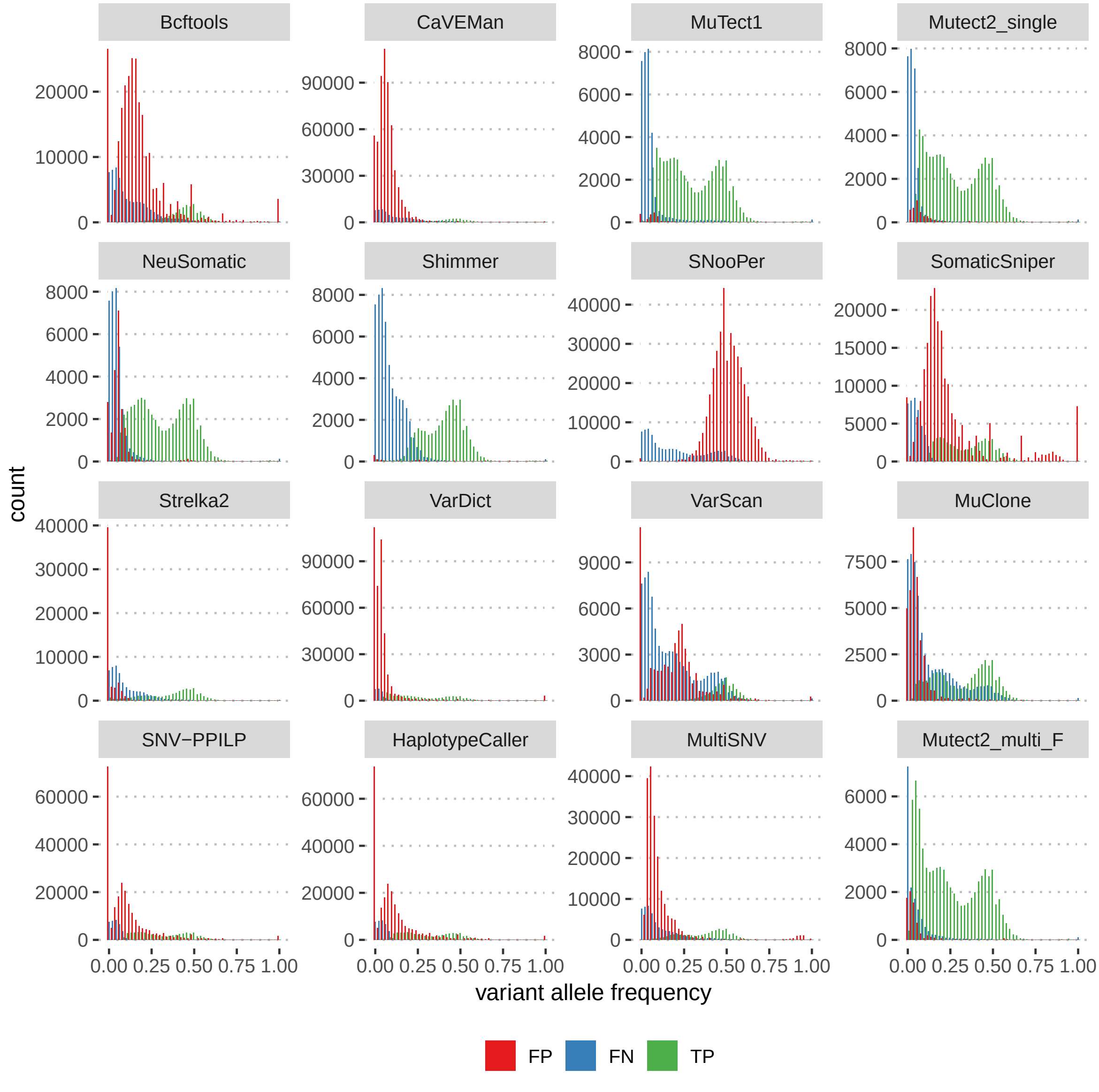


Figure S9: Variant allele frequencies in the *spike-in* simulation for true positive (TP), false positive (FP) and false negative (FN) calls. Y-axes were scaled individually for each subgraph.

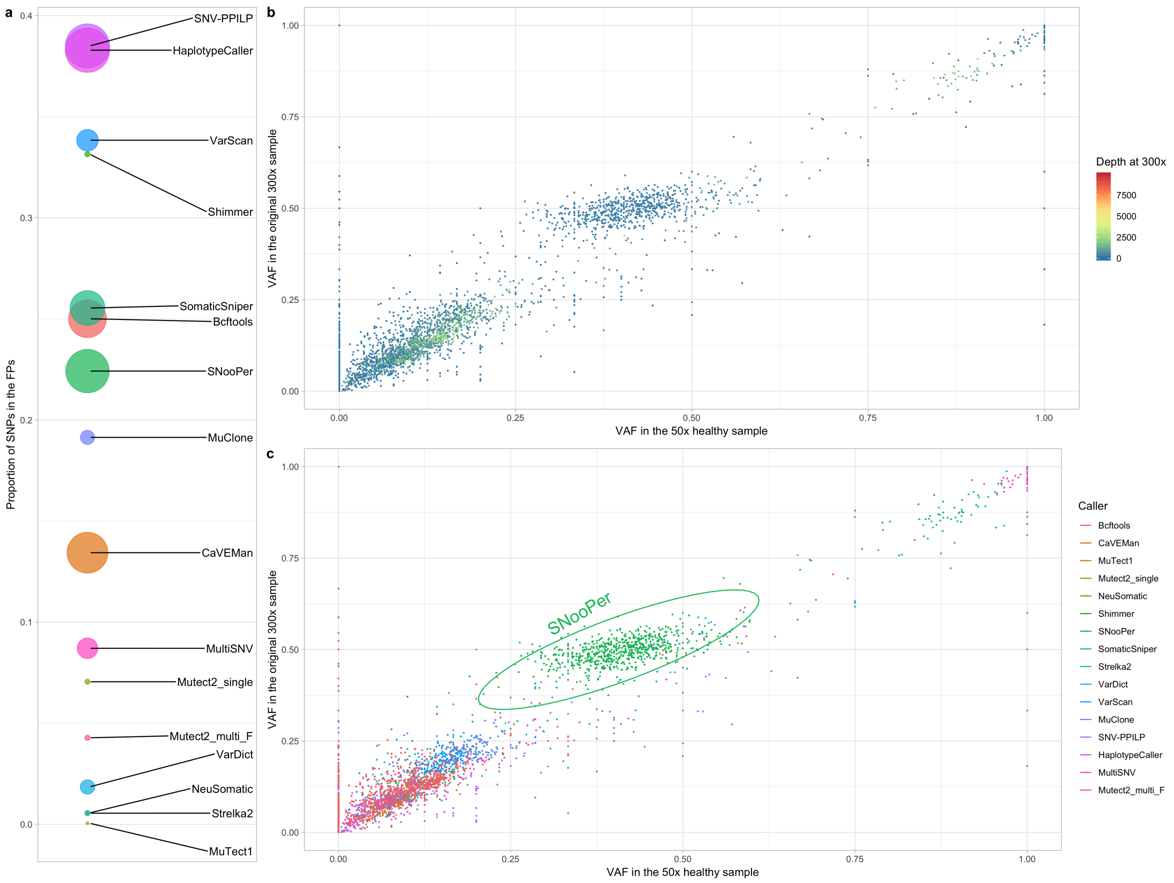


Figure S10: Germline variants detected as somatic variants in the *spike-in* simulations. **a** Percentage of FPs that correspond to SNPs for each caller. Dot size is proportional to the absolute number of FPs that correspond to SNPs. **b** VAF of the SNPs detected as somatic by any of the callers, both in the original 300x bam file (Y-axis) and in the downsampled 50x healthy used for the calling (X-axis). **c** VAFs in the original 300x bam file and in the downsampled 50x bam file of the SNPs called as somatic by each caller. Most germline variants present at frequencies close to 0.5 were called by SNooPer.

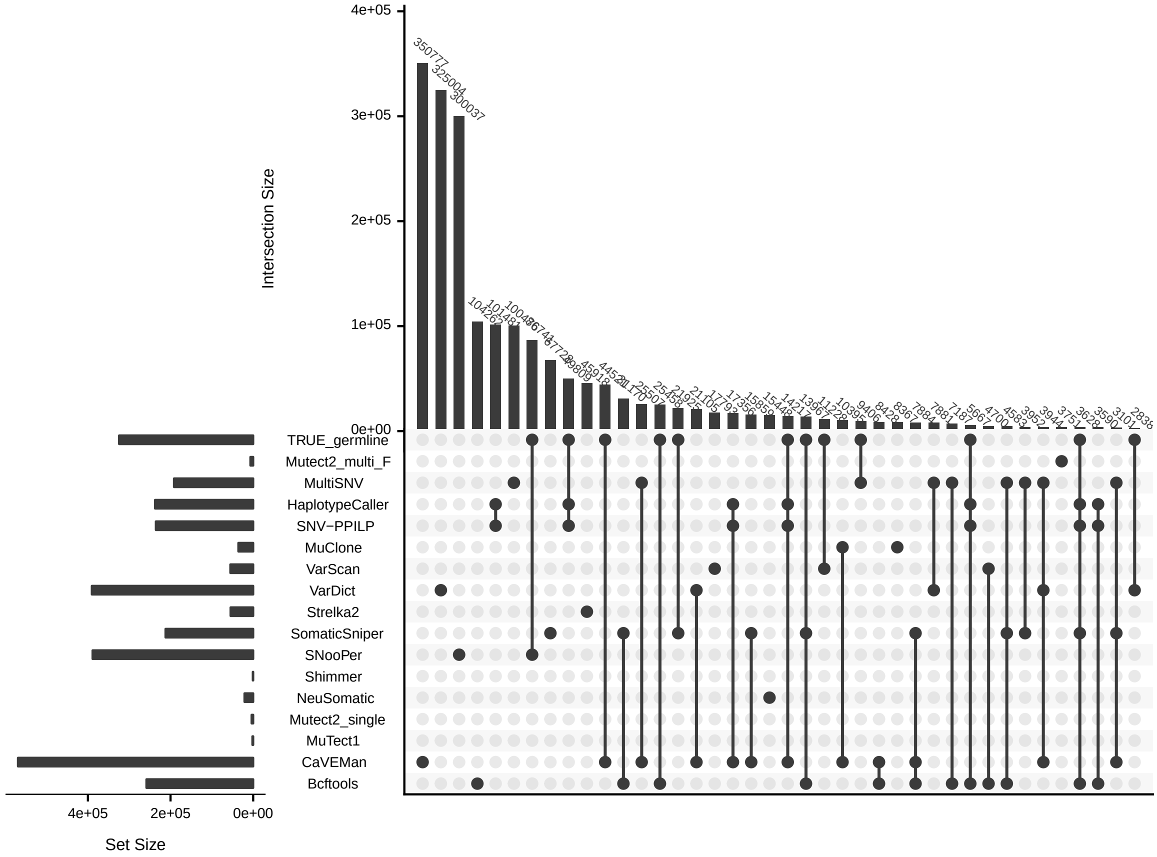


Figure S11: Intersection between false positive (FP) calls and germline variants in the *spike-in* simulations. TRUE_germline is the set of variants detected by HaplotypeCaller in the healthy sequencing data at 300x. Intersections involving TRUE_germline are false positives caused by SNPs. See Fig. S2 for a detailed explanation of the figure components.

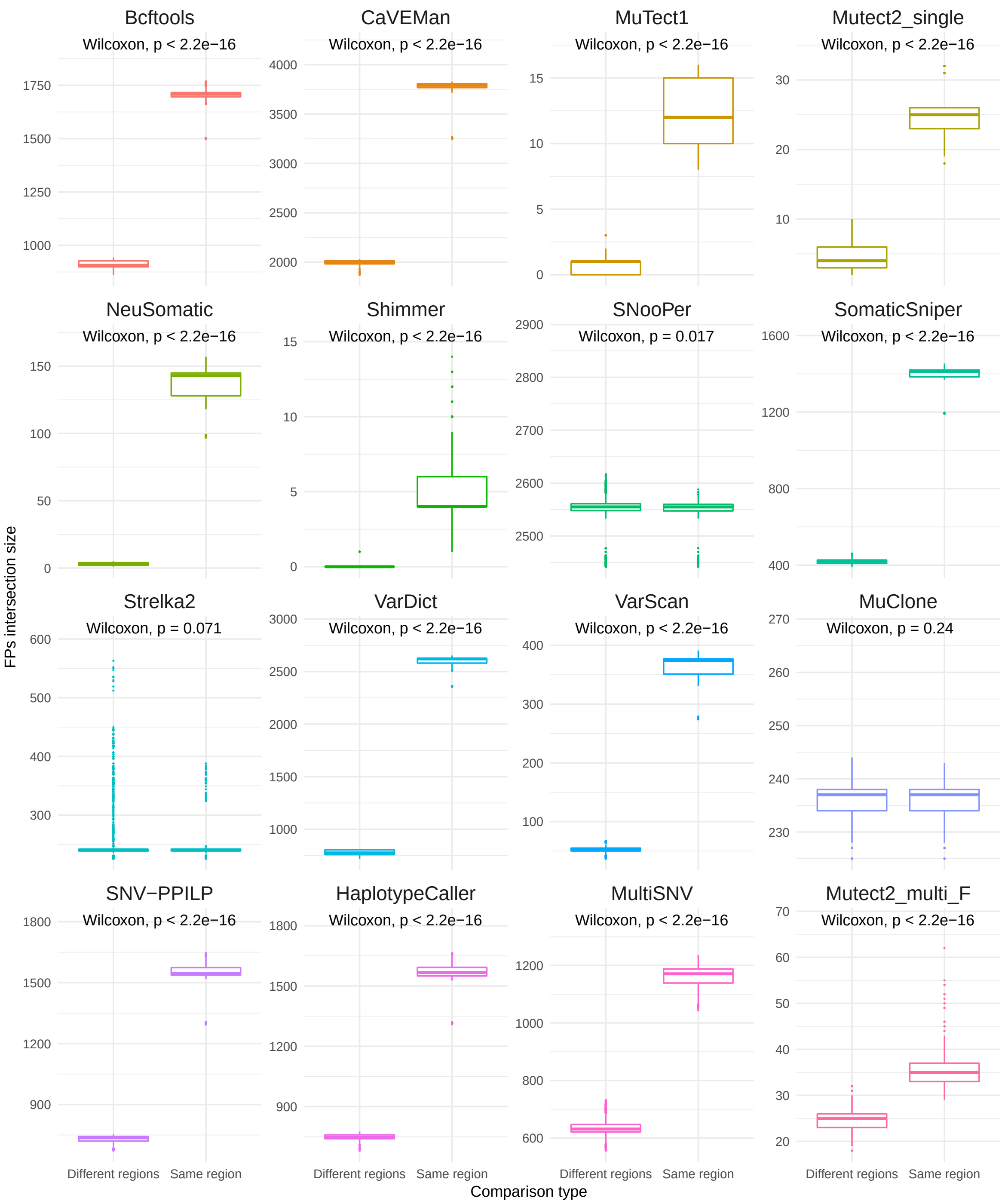


Figure S12: Recurrence of the false positives (FP) in the *spike-in* simulations. Pairwise FPs intersection between samples is shown for all the callers. Pairwise comparisons are split into comparisons between samples from the same region in different replicates, and different regions (in the same or in different replicates). The intersection between same-region samples is larger than the intersection between different-region samples. Boxplots: the central line indicates the median, while the box limits correspond to the $Q1$ and $Q3$ quartiles; upper and lower whiskers extend from $Q3$ to $Q3 + 1.5(Q3 - Q1)$ and from $Q1$ to $Q1 - 1.5(Q3 - Q1)$, respectively.

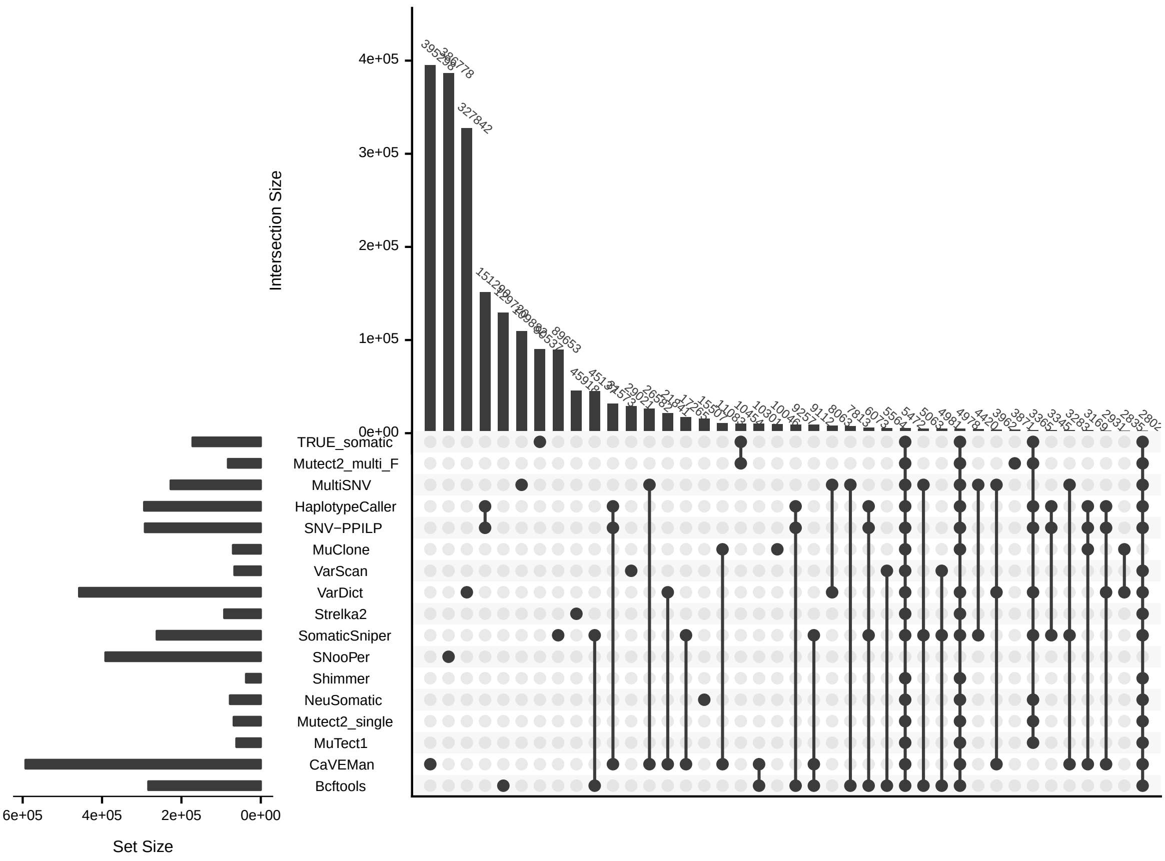


Figure S13: Intersection between variant call sets and true somatic variants in the *spike-in* simulations. TRUE_somatic is the set of spiked-in mutations. Intersections involving TRUE_somatic are true positives, while the rest are false positives. See Fig. S2 for a detailed explanation of the figure components.

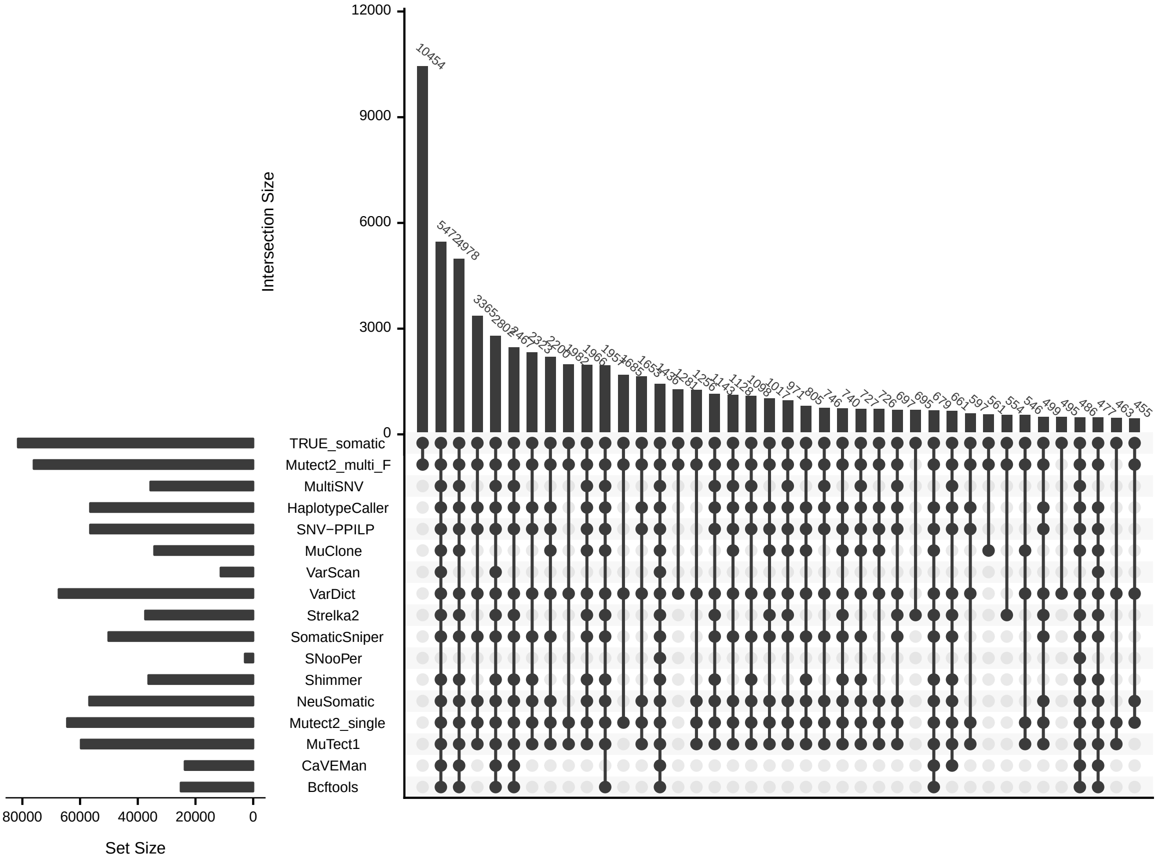


Figure S14: Intersection between true positive (TP) calls and true somatic variants in the *spike-in* simulations. TRUE_somatic is the set of spiked-in mutations. Only intersections involving TRUE_somatic (TPs) are shown. See Fig. S2 for a detailed explanation of the figure components.

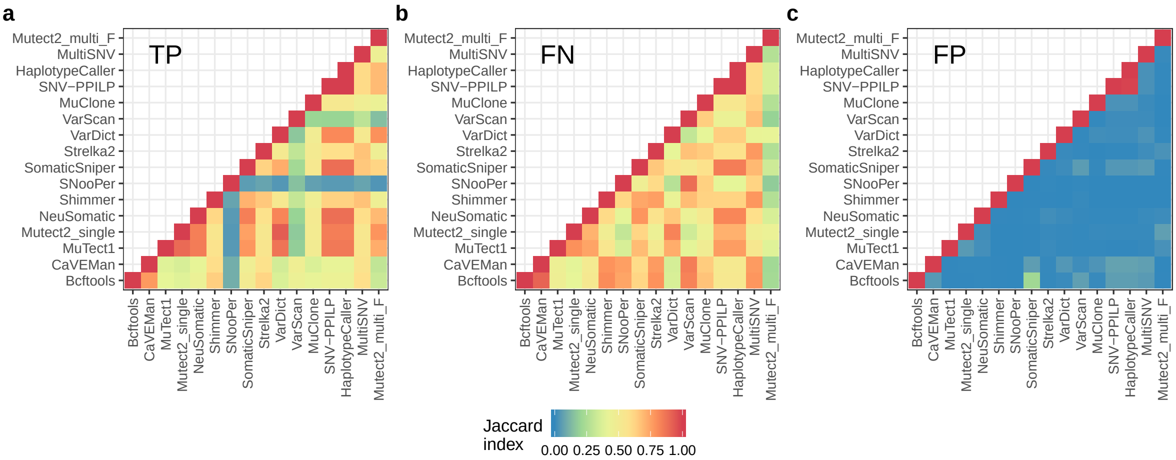


Figure S15: Jaccard distances between variant call sets in the *spike-in* simulations. **a** True positives (TP). **b** False negatives (FN). **c** False positives (FP).

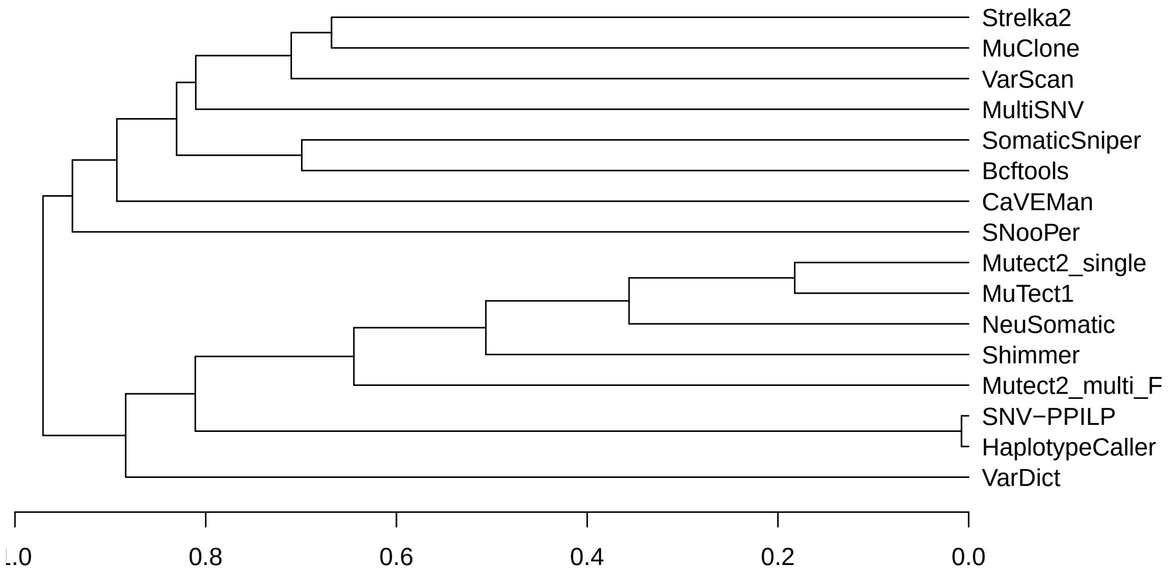


Figure S16: Hierarchical clustering of callers in the *spike-in* simulations. The clustering was carried out according to the Jaccard distance between the variant call sets for the different callers.

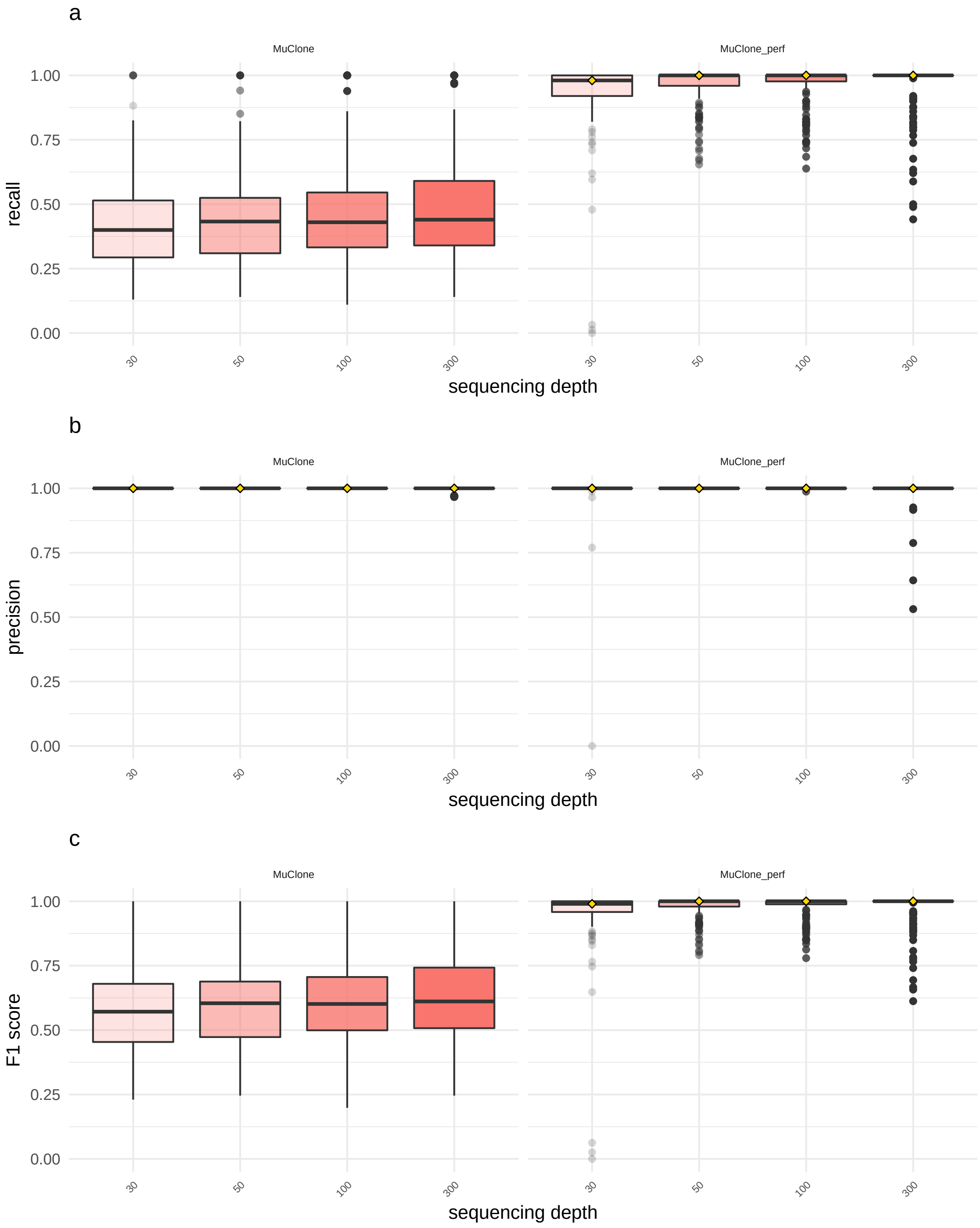


Figure S17: Performance of MuClone in the *de novo* simulations with and without clone prevalence information. The figure shows the recall (**a**), precision (**b**) and F1 score (**c**) at different sequencing depths of MuClone when the clonal prevalences were inferred (left, MuClone), and when the true clonal frequencies were used (right, MuClone_perf). For the former, filtered Mutect2_single variants were used to infer mutation clusters with PyClone; then, the unfiltered Mutect2_single variant set was classified by MuClone using the PyClone cluster frequencies as additional information. For the latter, ("perfect information") case, simulated variants plus the true clone frequencies were input to MuClone.
